# Multiplexed imaging of human tuberculosis granulomas uncovers immunoregulatory features conserved across tissue and blood

**DOI:** 10.1101/2020.06.08.140426

**Authors:** Erin F. McCaffrey, Michele Donato, Leeat Keren, Zhenghao Chen, Megan Fitzpatrick, Vladimir Jojic, Alea Delmastro, Noah F. Greenwald, Alex Baranski, William Graf, Marc Bosse, Pratista K. Ramdial, Erna Forgo, David Van Valen, Matt van de Rijn, Sean C. Bendall, Niaz Banaei, Adrie J.C. Steyn, Purvesh Khatri, Michael Angelo

**Affiliations:** Stanford University Department of Pathology; Stanford Center for Biomedical Informatics Research; Stanford Institute for Immunity, Transplantation and Infection; Calico Life Sciences LLC; University of Wisconsin, Department of Pathology; Division of Biology and Bioengineering, California Institute of Technology; African Health Research Institute; University of KwaZulu-Natal, Inkosi Albert Luthuli Central Hospital; Stanford University Department of Medicine-Infectious Diseases; Department of Microbiology, University of Alabama at Birmingham

## Abstract

Tuberculosis (TB) is an infectious disease caused by *Mycobacterium tuberculosis* that is distinctly characterized by granuloma formation within infected tissues. Granulomas are dynamic and organized immune cell aggregates that limit dissemination, but can also hinder bacterial clearance. Consequently, outcome in TB is influenced by how granuloma structure and composition shift the balance between these two functions. To date, our understanding of what factors drive granuloma function in humans is limited. With this in mind, we used Multiplexed Ion Beam Imaging by Time-of-Flight (MIBI-TOF) to profile 37 proteins in tissues from thirteen patients with active TB disease from the U.S. and South Africa. With this dataset, we constructed a comprehensive tissue atlas where the lineage, functional state, and spatial distribution of 19 unique cell subsets were mapped onto eight phenotypically-distinct granuloma microenvironments. This work revealed an immunosuppressed microenvironment specific to TB granulomas with spatially coordinated co-expression of IDO1 and PD-L1 by myeloid cells and proliferating regulatory T cells. Interestingly, this microenvironment lacked markers consistent with T-cell activation, supporting a myeloid-mediated mechanism of immune suppression. We observed similar trends in gene expression of immunoregulatory proteins in a confirmatory transcriptomic analysis of peripheral blood collected from over 1500 individuals with latent or active TB infection and healthy controls across 29 cohorts spanning 14 countries. Notably, PD-L1 gene expression was found to correlate with TB progression and treatment response, supporting its potential use as a blood-based biomarker. Taken together, this study serves as a framework for leveraging independent cohorts and complementary methodologies to understand how local and systemic immune responses are linked in human health and disease.

## Introduction

*Mycobacterium tuberculosis* (*Mtb*) is the leading cause of mortality from infectious disease in the world, accounting for nearly 1.5 million deaths each year^1^. Relative to other communicable diseases, the reduction in the incidence of TB infection over the last 20 years has been modest. This is largely due to the lack of a highly efficacious vaccine, lengthy and toxic antimicrobial regimens, and emergence of multidrug resistance. Along these lines, previous efforts to develop new host-directed therapies have been hindered by an incomplete understanding of how TB interacts with the human immune system during infection.

Infection is initiated when bacteria are engulfed by alveolar macrophages or other resident phagocytes after being inhaled into the lungs^2,3^. This triggers an immune response that ultimately converges on formation of a granuloma, a dynamic and spatially-organized tissue structure comprised of macrophages, granulocytes, lymphocytes, and fibroblasts. A prototypical non-necrotic granuloma consists of a myeloid-predominant central core region that is enriched with infected macrophages and encircled by lymphocytes. From the perspective of facilitating an effective host response, granulomas play seemingly contradictory roles. On the one hand, consolidation of infected cells within the myeloid core limits dissemination by partitioning them away from uninvolved lung parenchyma. On the other, tolerogenic pathways that are upregulated within this region limit bacterial clearance^4–6^.

Depending on the histological subtype and bacterial burden, granuloma composition can be highly variable^7^. This variability can manifest within a single individual, where infection can result in formation of multiple granulomas with distinct histologic and immunological features that each progress independently of one another over time^8^. Controlled infections in non-human primates (NHP) have revealed that a single individual can possess well over ten granulomas and that the inflammatory profile, size, and bacterial ecology of these lesions can vary dramatically^9–11^. Thus, the trajectory of each granuloma can vary across a spectrum between complete bacterial clearance to uncontrolled dissemination. This discordance suggests that local host-bacterial dynamics within the tissue microenvironment play a central role in determining granuloma fate. Along these lines, a growing number of studies suggest that granuloma structure and immune cell function are interconnected^12,13^. For example, previous work has suggested that impaired adaptive immunity might be the consequence of T cells being largely excluded from the myeloid-dominated central core region where infected macrophages tend to localize^14,15^.

Taken together, these findings suggest that TB disease progression is heavily impacted by focal, spatially-encoded regulatory mechanisms within the granuloma microenvironment. Consequently, understanding how these mechanisms promote bacterial clearance or persistence is critical for designing effective therapies that promote long term immunity. With this in mind, we used Multiplexed Ion Beam Imaging by Time-of-Flight (MIBI-TOF) to simultaneously image 37 proteins at subcellular resolution in solid tissue from individuals with active TB infection^16^. We compared granuloma composition with respect to 19 unique cell subsets to delineate different subtypes of granulomas that were enriched for classical monocytes, myeloid-derived suppressor cell-like (MDSC) macrophages, or tertiary lymphoid structures (TLS). We then utilized an adaptation of Latent Dirichlet Allocation (spatial-LDA) to identify spatially-coordinated immune responses within eight recurrent cellular microenvironments. These analyses revealed a microenvironment characterized by expression of regulatory proteins, IDO1 and PD-L1, and proliferative regulatory T cells. Paradoxically, these cells were not accompanied by significant numbers of PD-1^+^ lymphocytes or any other evidence suggesting T cell exhaustion.

To determine if these features were associated with drug treatment or severity of infection, we leveraged publicly available gene expression data of peripheral blood from patients with TB. In line with the granuloma imaging data, we found increased expression of *IDO1* and *CD274* (PD-L1) and diminished expression of *PDCD1* (PD-1) and *LAG3* in individuals with active TB. Moreover, *CD274* (PD-L1) gene expression was found to associate with progression from latent to active disease and with therapeutic response, suggesting it could be used as a novel prognostic biomarker. Taken together, this work provides compelling evidence for a myeloid-mediated mechanism of immune suppression driven locally within the granuloma that promotes bacterial persistence and subverts T-cell activation.

## Results

### Multiplexed imaging of human tuberculosis granulomas reveals structured immune cell composition

To assess granuloma composition and architecture in TB, we curated a cohort of actively infected human tissues. Archival formalin-fixed paraffin-embedded (FFPE) specimens from patients treated in the United States or South Africa were procured from Stanford Health Care or University of KwaZulu-Natal, Inkosi Albert Luthuli Central Hospital, respectively (Extended Data Table 1). The South African samples were pulmonary tissues from patients that underwent therapeutic resection for advanced TB (n = 6). While TB disease typically manifests in the lung, infection can disseminate to extra-pulmonary sites as well^17,18^. To characterize TB infection at an earlier stage and assess how granuloma composition varies with infection site, samples from US patients consisted of diagnostic biopsies from lung (n = 2), pleural cavity (n = 3), lymph node (n = 1), and endometrium (n = 1) (Fig. 1a).

**Figure 1.**
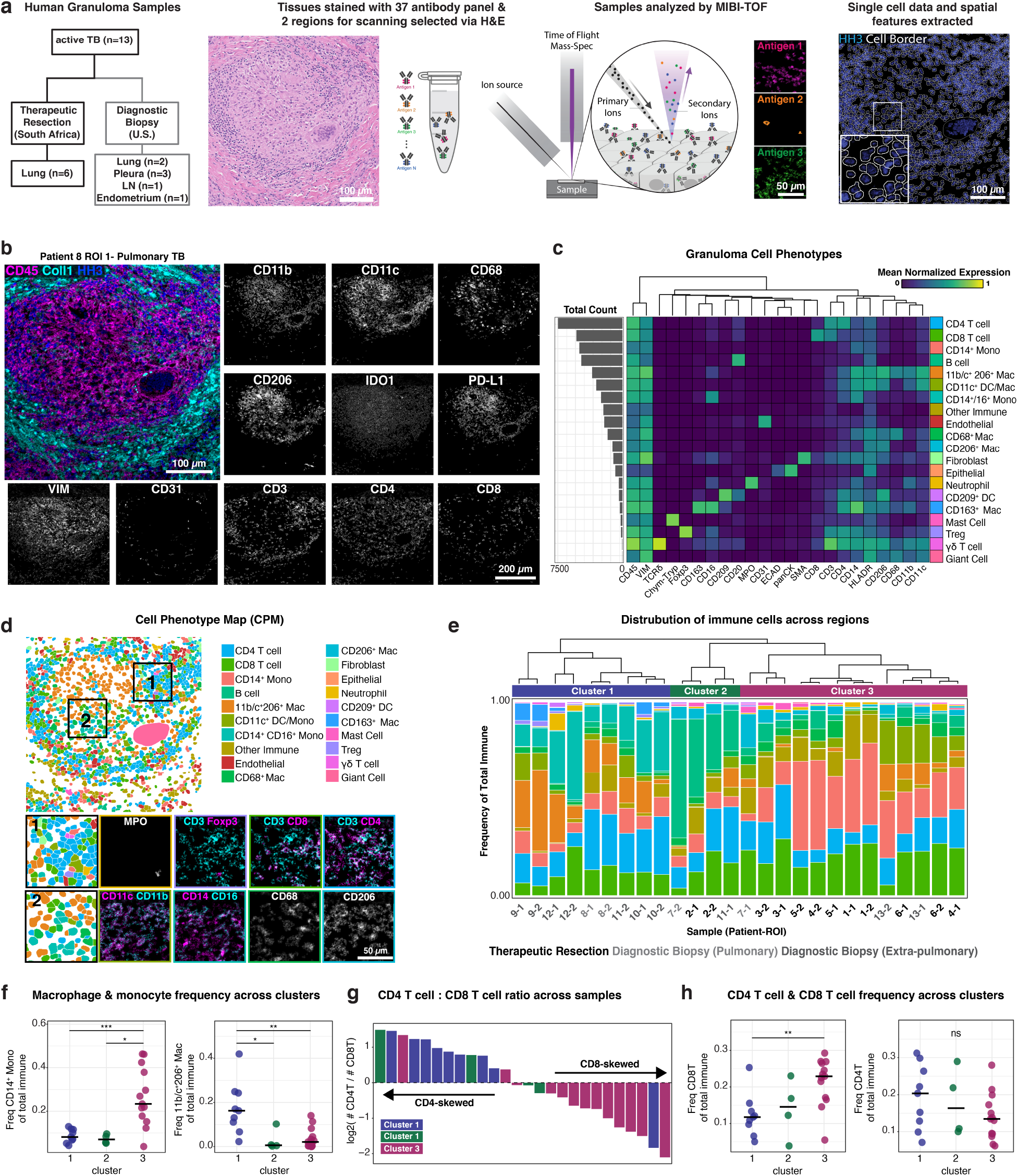
Multiplexed imaging of human tuberculosis granulomas reveals structured immune cell composition. **(a)** Conceptual overview of MIBI-TOF analysis of human TB granulomas. **(b)** Representative images from a pulmonary TB section. **(c)** Cell lineage assignments based on normalized expression of lineage markers (heatmap columns). Rows are ordered by absolute abundance shown on the bar plot (left), while columns are hierarchically clustered (Euclidean distance, average linkage). **(d)** Cell identity overlaid onto the segmentation mask for a representative pulmonary TB section (left). Two insets (bottom) are shown. **(e)** The relative abundance of immune cell types across all TB FOVs with cell types ordered by decreasing median abundance and bars ordered by the hierarchical relationship of pairwise Pearson correlation coefficients (distance = 1 – correlation, complete linkage). Consensus clusters are annotated above bar plot (cluster 1 = blue, cluster 2 = green, cluster 3 = purple). **(f)** Frequency of CD14+ monocytes and 11b/c^+^ 206^+^ macrophages of total immune cells colored by cluster. Line represents the median. **(g)** The CD4 T: CD8 T cell ratio represented as a log2 fold-change for each TB FOV (top) colored by cluster. **(h)** Frequency of CD4^+^ and CD8^+^ T cells of total immune cells colored by cluster. Line represents the median. All p-values determined with a Wilcoxon Rank Sum Test where: ns p > 0.05, * p < 0.05, ** p < 0.01, *** p < 0.001.

Tissue sections for each specimen were reviewed by an anatomic pathologist and screened to include the presence of solid, non-necrotic granulomas or active granulomatous inflammation, while excluding excessively necrotic or fibrotic regions (Extended Data Fig. 1a). MIBI-TOF was subsequently used to image two 500 μm^2^ fields of view (FOVs) per tissue stained with a 37-plex panel of metal-labeled antibodies (Fig. 1b, Extended Data Fig. 1b-c, Extended Data Table 2)^16^. The antibody panel included markers to comprehensively phenotype most major immune and non-immune cell lineages, including lymphocytes, macrophages, granulocytes, stroma, and epithelium. The panel also included antibodies for 12 functional markers with an emphasis on those with well-documented immunoregulatory pathways, including PD-1, Lag3, PD-L1, and IDO1.

To extract single cell phenotypes, multiplexed imaging data were processed with a low-level pipeline prior to single-cell segmentation (Fig. 1a, Extended Data Fig. 1d)^19–21^. Each FOV contained an average of ~1400 single cells (sd = 312) (Extended Data Fig. 2d). Hierarchical application of the FlowSOM algorithm (Extended Data Fig. 2a, b) was employed to phenotype 19 unique cell subsets (Fig. 1c)^22^ using cell area normalized values of protein expression for each segmented cell event. For each image, FlowSOM clusters and segmentation masks were combined to generate cell phenotype maps (CPM) where each cell is labeled by its phenotype (Fig. 1d, Extended Data Fig. 2c).

Consistent with previous work, granuloma composition was predominated in most lesions by T cells and myeloid cells, (average myeloid: lymphoid ratio = 1.4, sd = 1.0)^23^. Myelomonocytic cells were comprised of multiple subsets of macrophages, dendritic cells, and monocytes that were each distinguished by varying degrees of co-expression of CD11c, CD11b, CD209, CD68, CD14, CD16, and CD206 (Fig. 1e). Granulocytes were comprised largely of neutrophils (mean = 1.2%, sd = 1.7) and mast cells (0.6% ± 0.8)^24–26^. We also identified γδ T cells (0.1% ± 0.22), CD209^+^ dendritic cells (0.2% ± 0.5), and regulatory T cells (0.4% ± 0.6), highlighting the capability of our approach to enumerate low abundance immune cell populations that have been suggested to play a key role in granuloma pathology. In line with reports of increased vascularization in active disease, non-immune cells were predominated by endothelial cells (5.7% ± 3.1) while fibroblasts (3.3% ±5.1) and epithelial cells (2.7% ± 4.0) varied significantly between lesions (Extended Data Fig. 2e, f)^27,28^. Altogether, we assigned 94% (n = 33,194 single cells) of cells to 19 subsets that in aggregate ranged in frequency from 0.1-15% across our dataset.

To assess whether granulomas can be grouped into subtypes based exclusively on immune cell subset frequencies, we clustered FOVs using Pearson correlation based on their immune cell proportions (Extended Data Fig. 2g). We found that using three clusters explained 72% of the variance in our dataset (Fig. 1e, Extended Data Fig. 2h). Cluster 1 (Fig. 1e, f) was characterized by CD11c^+^ CD11b^+^ macrophages, intermediate monocytes, and occasional CD163^+^ macrophages. Chi-square analysis of cell type co-occurrence showed significant associations of CD163^+^ macrophages with intermediate monocytes (adj. p = 0.0090), regulatory T cells (Tregs) (adj. p = 0.003), and fibroblasts (adj. p = 0.02), suggesting coordinated cellular presence in this cluster (Extended Data Fig. 2i). Cluster 2 was enriched for B cells (Fig. 1e). Lastly, a subset of granulomas (cluster 3) was enriched for classical monocytes (Fig. 1e, f) and higher numbers of CD8^+^ T cells, resulting in a skewed CD4^+^ to CD8^+^ T cell ratio relative to clusters 1 and 2. Since most samples in cluster 3 were surgical lobectomies from patients in South Africa, this profile could be related to disease severity, comorbidity, or mandatory pre-surgical antimicrobial therapy. Given this result, we analyzed therapeutic resections with respect to HIV status (Extended Data Fig. 3). While we observed a slight difference in the CD4^+^ to CD8^+^ T cell ratio (approximately two-fold decrease in HIV^+^ specimens, p = 0.026), we found no material differences in immune cell frequencies that define these clusters. Taken together, this comprehensive cell census reveals distinct types of granulomas that are defined by immune cell frequency and associate with TB disease status.

### Spatial analysis of granulomas identifies organized protein expression patterns and conserved cellular microenvironments

In order to identify recurrent patterns of protein co-expression in TB granulomas, we conducted a spatial enrichment analysis that quantified the degree of co-occurrence between pairs of proteins relative to a null distribution (Extended Data Fig. 4a)^19^. Pairwise enrichment scores for each protein pair were then used to construct an interaction network that was subsequently analyzed using a community detection algorithm^29^ (Fig. 2a). This analysis resulted in three spatial modules consistent with canonical granuloma structures, including the myeloid core, lymphocytic cuff, and stromal compartment. Intriguingly, these modules also revealed more granular, previously unknown features linking cell function to spatial organization, such as the association of the lymphocytic cuff with H3K9Ac and the myeloid core with IDO1 and PD-L1 (Fig. 2a).

**Figure 2.**
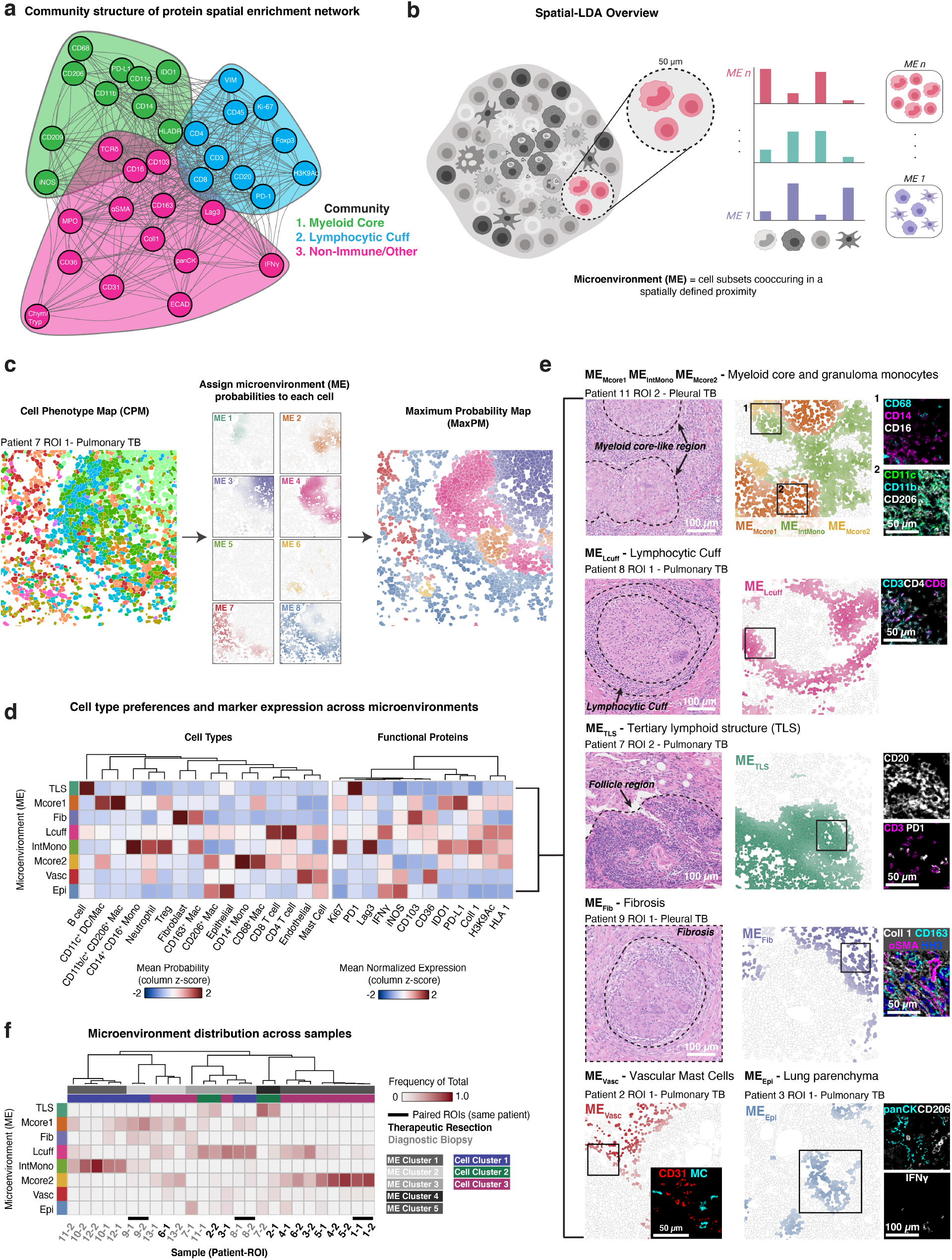
Spatial analysis of granulomas identifies organized protein expression patterns and conserved cellular microenvironments. **(a)** Positive spatial enrichments (average z-score > 0) between protein pairs as a weighted, undirected network (edge weight is proportional to average z-score) with three communities (myeloid core = green, lymphocytic cuff = blue, non-immune/other = pink). **(b)** Conceptual overview of spatial-LDA. **(c)** Cell probability map (left), max probability map (right), and microenvironment (ME) probability for 8 MEs (middle, scaled 0 −1) for a pulmonary TB section. **(d)** Heatmap of ME preferences for all subsets (standardized mean ME loading) with hierarchical clustering (Euclidean distance, complete linkage) and mean normalized expression of functional markers (probability weighted mean) with columns hierarchically clustered (Euclidean distance, complete linkage). **(e)** Biological classification of microenvironments. **(f)** Frequency of all MEs per FOV. Heatmap columns are hierarchically clustered (Pearson correlation, complete linkage). Paired ROIs from the same patient annotated with a black bar. ME and cell composition clusters annotated below dendrogram. Cell cluster annotation is based on clusters in Fig. 1e.

These findings motivated us to examine how the local cellular neighborhood relates to single cell function and granuloma structure. Therefore, we employed spatial Latent Dirichlet Allocation (spatial-LDA)^30^ to discover and assign cellular microenvironments (MEs) to each cell in a CPM, where an ME is defined by a set of cell types found to be spatially co-occurring across the cohort. Using this approach, we identified eight MEs for summarizing the local frequency of cell subsets within a 50 μm radius of a target cell (Fig. 2b). We then labeled each cell with its highest probability ME to generate a maximum probability ME map (MaxPM, Fig. 2c, Extended Data Fig. 4b). Through this approach, granuloma composition and structure can be summarized with two complementary and simplified spatial representations: a CPM and MaxPM where cells are labeled either by cell type or by ME, respectively (Fig. 2c).

This allowed us to annotate well known features of granuloma histology in an unbiased fashion, while also revealing previously unrecognized cellular niches (Fig. 2d, e). For example, the large majority of granuloma macrophages and monocytes were assigned to one of three myeloid MEs (ME_Mcore1_, ME_IntMono_, ME_Mcore2_). While MEMcore1 was found to some degree across all specimen types, ME_IntMono_ and ME_Mcore2_ were significantly enriched in either extra-pulmonary diagnostic biopsies or therapeutic resections, respectively (Fig. 2f, Extended Data Fig. 4d, e). ME_Mcore1_ exhibited the strongest preference for the histologically defined granuloma core region (median frequency in core = 99.1%, Extended Data Fig. 4c) and was predominated by CD11c^+^ CD11b^+^ macrophages (Fig. 2d, e). ME_Mcore2_ exhibited myeloid core preference to a lesser extent and was enriched for CD14^+^ classical monocytes (median frequency in core = 60.9%, Extended Data Fig. 4c). Lastly, ME_IntMono_ exhibited low myeloid core preference and was enriched for CD14^+^ CD16^+^ intermediate monocytes (Fig. 2d, e).

ME_Lcuff_ aligned with the second histologically defined microenvironment in the granuloma, the lymphocytic cuff, and was comprised predominantly of CD4^+^ and CD8^+^ T cells (Fig. 2d, e). METLS is a second lymphoid ME that is B cell predominated with sparse numbers of follicular helper T cells (CD4^+^ PD-1^+^), consistent with tertiary lymphoid structures (TLS, confirmed by H&E in serial sections, Fig. 2e)^31–33^. This ME was highly abundant in FOVs that were B cell predominated from cell frequency cluster 2 (Fig. 2f, Fig. 1e).

As previously demonstrated, some granulomas exhibited a fibrotic wound healing response consisting of fibroblasts and CD163^+^ M2-like macrophages (Fig. 1e). These cells were found to co-localize within ME_Fib_, where CD36, a fibroblast marker, and collagen-1, a marker for fibrosis are expressed (Fig. 2d, e)^34^. The last two MEs represented less characterized cellular environments in TB infection. ME_Vasc_ was predominated by blood vessels and mast cells while ME_Epi_ was comprised of parenchymal epithelial cells and CD206^+^ alveolar-like macrophages (Fig. 2d, e). Given that they are known to participate in angiogenesis, tissue repair, and immune cell recruitment^35^, perivascular localization of mast cells in the granuloma could suggest their involvement in some of these processes. On the other hand, since ME_Vasc_ was found to be lower in extra-pulmonary biopsies (Fig. 2f, Extended Data Fig. 4e), this may reflect organ-specific differences in vascularity and abundance of tissue resident mast cells.

We next sought to compare sample composition with respect to ME frequency. Using a correlation-based approach, we found that five ME frequency clusters accounted for 89% of variance (Fig. 2f, Extended Data Fig. 4f). Notably, two of these clusters were comprised of samples from more than one cell frequency cluster (as defined in Fig. 1e). These clusters, along with a Principle Component Analysis (PCA) of all samples based on mean ME probability, further supported tissue site and clinical cohort-associated ME profiles (Fig. 2f, Extended Data Fig. 4g). Altogether, this suggests that MEs capture recurrent spatial features of granulomas that are not discernible by bulk cell composition alone.

### Granuloma myeloid cells are characterized by spatially coordinated expression of IDO1 and PD-L1

Our microenvironment and spatial protein network modeling revealed that myeloid-rich regions in the granuloma are characterized by expression of two proteins, IDO1 and PD-L1 (Fig. 2a, d). Given the tolerogenic role of these proteins^36–41^, we sought to identify myeloid cell subsets that could promote bacterial persistence through upregulation of these pathways. We identified nine unique macrophage, monocyte, and dendritic cell populations (Extended Data Fig. 2b, Fig. 3a). PD-L1 and IDO1 were correlated (Pearson R = 0.64, p < 2.2 x 10^-16^) and expressed to varying degrees across most of these populations and all FOVs (Fig. 3b-e, Extended Data Fig. 5a, c). Bright co-expression of both proteins was observed in CD11b^+^ CD11c^+^ macrophages, consistent with an MDSC-like phenotype^42^ (Fig. 3d). This was also found in CD209^+^ DCs and CD16^+^ CD14^+^ intermediate monocytes, where it was associated with HLA-DR downregulation (Fig. 3a-d). While the frequency of IDO1^+^ cells did not vary significantly between tissue or specimen type (Extended Data Fig. 5b), PD-L1 ^+^ cells were significantly higher in diagnostic biopsies relative to therapeutic resections, with a notable enrichment in extrapulmonary tissues (Extended Data Fig. 5b). Notably, neutrophils were also found to express IDO1 or PD-L1 (Extended Data Fig. 5d). Given that they have been shown to secrete anti-inflammatory cytokines in TB granulomas^43^, these findings are consistent with a regulatory effector function. Lastly, nearly 100% of multinucleated giant cells expressed IDO1 and ~75% express PD-L1 (Fig. 3f).

**Figure 3.**
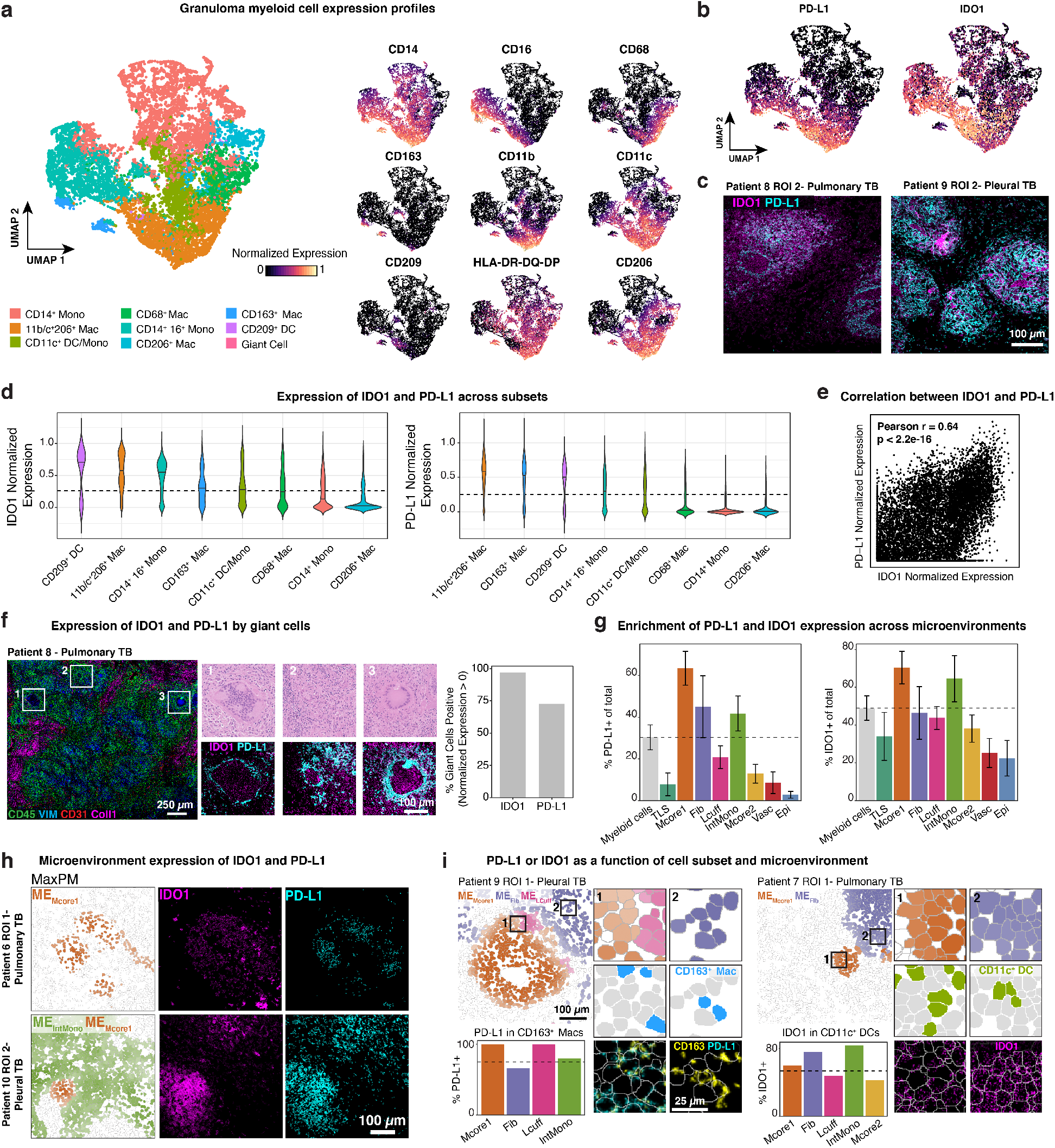
TB granuloma myeloid cells are characterized by spatially coordinated expression of IDO1 and PD-L1. **(a)** UMAP visualization of all myeloid populations across all TB FOVs colored by subset (left) and normalized expression of phenotypic markers used to delineate subsets. **(b)** IDO1 and PD-L1 normalized expression overlaid on the UMAP. **(c)** Representative images of a pulmonary (left) and pleural (right) sample showing expression of IDO1 (magenta) and PD-L1 (cyan). **(d)** Normalized expression of IDO1 (top) and PD-L1 (bottom) for major myeloid subsets ordered by decreasing median expression value. Dashed line indicates the cutoff for positivity for IDO1 (cutoff = 0.26) and PD-L1 (cutoff = 0.25). **(e)** PD-L1 and IDO1 expression values across all myeloid cells as a biaxial scatter plot. Plot displays Pearson’s r and p-value determined by t-test. **(f)** Giant cells identified from a MIBI-scanned pulmonary TB sample (CD45 = green, Vimentin = blue, CD31 = red). Representative giant cells displayed in zoomed insets (lower left) with hematoxylin and eosin staining or IDO1 (magenta) and PD-L1 (cyan) expression. Bar plot displays the percentage of IDO1+ and PD-L1^+^ giant cells (n = 33, normalized expression > 0). **(g)** ME_Mcore1_ and ME_IntMono_ max probability maps and representative images of a pulmonary (top) and pleural (bottom) TB sample showing expression of IDO1 (magenta) and PD-L1 (cyan). **(h)** The frequency of IDO1^+^ and PD-L1^+^ myeloid cells for all myeloid cells in aggregate and broken down by microenvironment. **(i)** The frequency of PD-L1^+^ CD163^+^ macrophages (left) across MEs with a representative MaxPM. Insets are colored by ME (top), cell type (blue, middle), and CD163 (yellow) and PD-L1 (cyan) with the segmentation boundaries overlaid (white). The frequency of IDO1 ^+^ CD11b^+^ CD11c^+^ macrophages (right) across MEs with a representative MaxPM. Insets are colored by ME (top), cell type (green, middle), and IDO1 (magenta) with the segmentation boundaries overlaid (white).

To determine how PD-L1 and IDO1 expression is associated with a cell’s location in the granuloma, we calculated the frequency of PD-L1^+^ and IDO1^+^ cells for each cell subset in each ME (Fig. 3i, Extended Data Fig. 5e). We found that the majority of cells displayed preferential, ME-specific expression patterns. For example, the frequency of PD-L1 expressing CD163^+^ macrophages was highest in the ME_Mcore1_ (100%) and ME_Lcuff_ (100%), while IDO1 expressing CD11b^+^ CD11c^+^ macrophages were most enriched in ME_Fib_ (75.6%) and ME_IntMono_ (83.3%) (Fig. 3i). Altogether, PD-L1 and IDO1 expression defines a newly identified, spatially-coordinated immunoregulatory feature of TB granulomas that supports the possibility of highly localized, myeloid-mediated immune suppression in the granuloma.

### Granuloma lymphocytes are enriched in distinct cellular microenvironments and display a paradoxical absence of exhaustion markers

We next wanted to assess if the spatial coordination observed in tolerogenic myeloid cells extended to tolerogenic lymphocytes, like regulatory T cells (Tregs), which comprised 1.3% of all lymphocytes (Fig. 4a). Tregs (CD3^+^ CD4^+^ Foxp3^+^) were preferentially enriched in ME_Mcore1_ relative to ME_Lcuff_ (Fig. 4b, p = 0.0012), which stood in contrast to all other lymphocyte subsets, including Foxp3^−^CD4^+^ T cells. Furthermore, the frequency of proliferating Tregs exceeded that of all other major lymphocyte subsets (Fig. 4c, p < 0.001). These results suggest that Tregs and MDSC-like cells preferentially colocalize within ME_Mcore1_ to potentiate an immunomodulatory niche that could ultimately deter bacterial clearance (Fig. 4d)^44–50^.

**Figure 4.**
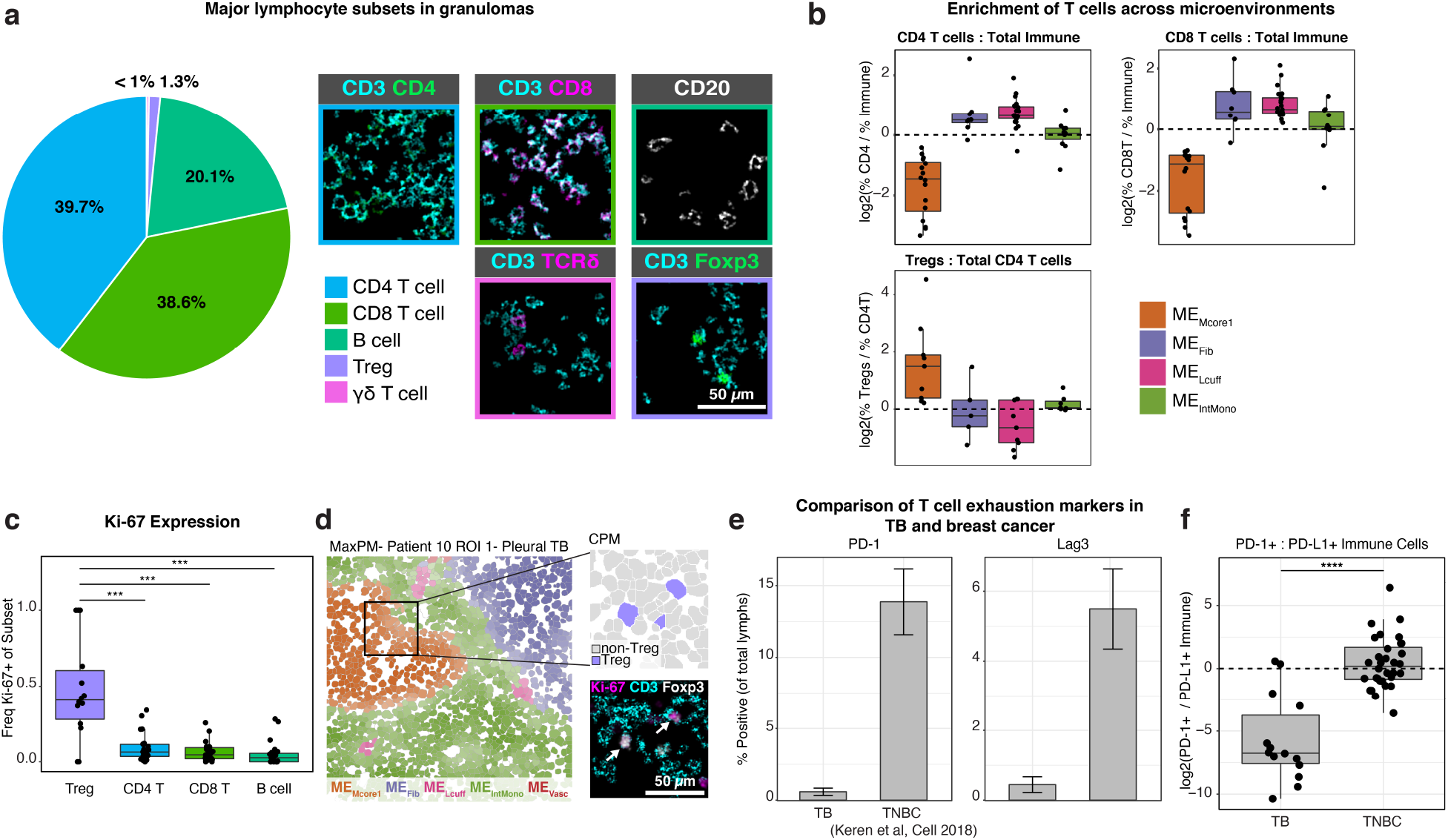
Granuloma lymphocytes are enriched in distinct cellular microenvironments and display a paradoxical absence of exhaustion markers. **(a)** Frequency of lymphocyte subsets in all TB FOVs pooled together (left) and representative images of each subset (right). **(b)** The frequency of CD4^+^ and CD8^+^ T cells relative to the frequency of total immune cells in four MEs of interest (top). The frequency of Tregs relative to the frequency of total CD4^+^ T cells (lower, left). **(c)** Frequency of Ki-67^+^ cells broken down by lymphocyte subset. **(d)** Representative image from a pleural TB FOV, colored by ME assignment (left). Zoomed inset displays Treg assignment (upper-right, purple = Treg, grey = non-Treg) and expression of Ki-67 (magenta), CD3 (cyan), and Foxp3 (white) (lower-right). **(e)** Percent of lymphocytes positive for PD-1 (left) and Lag3 (right) in all TB FOVs and TNBC. Bar represents the mean and standard error. **(f)** The ratio of PD-1^+^: PD-L1^+^ immune cells represented as a log2 fold-change in all TB FOVs and TNBC. All boxplots represent the median and interquartile range. All p-values determined with a Wilcoxon Rank Sum Test where: ns p > 0.05, * p < 0.05, ** p < 0.01, *** p < 0.001, **** p < 0.0001.

Anti-inflammatory pathways like those found in ME_Mcore1_ are often induced as a form of negative feedback that moderates the cytotoxic effects of unchecked immune activation^51^. In line with this, high expression of PD-L1 and IDO1 by granuloma myeloid cells would be expected to be accompanied by evidence of T cell activation in the form of checkpoint marker expression (e.g. PD-1, Lag3)^52^. For example, in previous work examining infiltrated triple negative breast cancer (TNBC) tumors, we found the median ratio of PD-1^+^: PD-L1^+^ immune cells to be near unity (Fig. 4f) and the prevalence of Lag3 or PD-1 positive lymphocytes to be 13.9% and 5.5% on average, respectively (Fig. 4e). In contrast, PD-L1 ^+^ granuloma myeloid cells far outnumbered PD-1 ^+^ lymphocytes (log2[PD-1 ^+^: PD-L1 ^+^] = −5.73 ± 3.4, Fig. 4f). Furthermore, the small numbers of PD-1 ^+^ lymphocytes in our dataset were almost entirely restricted to METLS, consistent with T follicular helper cells rather than an activated or exhausted phenotype (Extended Data Fig. 5f). These findings are consistent with reports from the cynomolgus macaque model that also found low levels of PD-1, Lag3, and CTLA4^14^, suggesting that PD-L1 expression by myeloid granuloma cells occurs independently of local cytokine release by activated T cells.

### TB and sarcoidosis granulomas have both common and diverging features of immune regulation

In addition to being the histological hallmark of TB, granulomatous inflammation can occur in response to a foreign body or in autoimmune disorders, such as sarcoidosis^53^. Interestingly, several gene expression studies that have attempted to develop blood-based biomarkers for latent and active infection have struggled to differentiate between TB and sarcoidosis^54,55^. In order to determine to what extent the features identified here overlap with other granulomatous diseases, we compared the TB sample cohort to ten sarcoidosis cases (Extended Date Fig. 6a-b). TB lesions were more variable in composition (p = 0.037, Extended Date Fig. 6d) and had significantly higher frequencies of CD8^+^ T cells, neutrophils, intermediate monocytes, and giant cells (Fig. 5a, Extended Data Fig. 6b-c). Sarcoid granulomas were heavily CD4^+^ T cell-skewed, even relative to the CD4-skewed TB granulomas in our dataset, consistent with reports of the pathology being driven primarily by Th17 and Th1 T cells (Fig. 5b)^56–58^.

**Figure 5.**
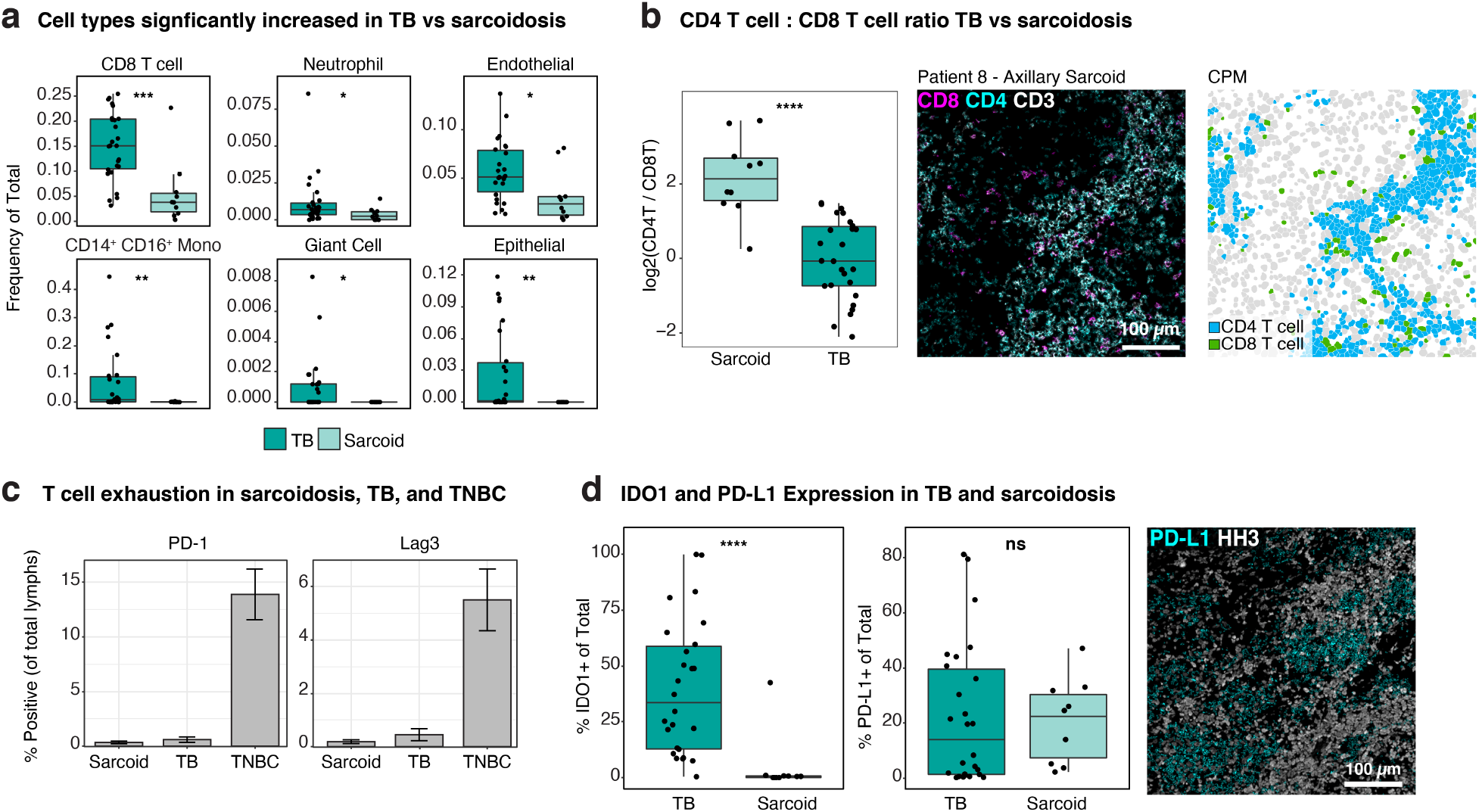
TB and sarcoidosis granulomas have both common and diverging features of immune regulation. **(a)** Frequency of cell subsets (of total cells) in TB versus sarcoidosis that are significantly different. **(b)** Comparison of the CD4^+^ T: CD8^+^ T cell ratio in TB versus sarcoidosis. Representative image of an axillary sarcoidosis FOV showing expression of CD8 (magenta), CD4 (cyan), and CD3 (white) (left) and colored by cell type (right, blue = CD4^+^ T cell, green = CD8^+^ T cell) **(c)** Percent of lymphocytes positive for PD-1 (left) and Lag3 (right) in all sarcoidosis FOVs, TB FOVs and TNBC. Bar represents the mean and standard error. **(d)** Percent of total cells positive for IDO1 or PD-L1 in TB and sarcoidosis. Representative image of a sarcoidosis FOV showing expression of PD-L1 (cyan) and HH3 (white). All boxplots represent the median and interquartile range. All p-values determined with a Wilcoxon Rank Sum Test where: ns p > 0.05, * p < 0.05, ** p < 0.01, *** p < 0.001, **** p < 0.0001.

Like TB, sarcoid lesions were PD-1 and Lag3 depleted (Fig. 5c) despite high levels of PD-L1^+^ myeloid cells (Fig. 5d, Extended Data Fig. 6e). However, unlike TB, IDO1 expression in sarcoid samples was almost entirely absent (Fig. 5d). Since we used a conservative threshold for IDO1 and PD-L1 positivity, our analysis biased toward the moderate to bright-expressing cells present in TB granulomas and control tissues. Therefore, to more comprehensively evaluate the disease specificity of PD-L1 and IDO1, we performed immunohistochemistry (IHC) for both proteins on a tissue microarray of granulomas from sarcoidosis (n = 9), foreign body uptake (n = 4), endometriosis (n = 4), and xanthomatosis (n = 3) (Extended Data Fig. 6f). We identified weak expression of IDO1 in several sarcoidosis lesions along with bright expression of PD-L1 as observed by MIBI-TOF (Extended Data Fig. 6f). However, IDO1^+^ and PD-L1^+^ cells were nearly absent in all xanthomas and endometrial lesions and rare in foreign body granulomas.

Notably, we observed similarly high levels of IDO1 and PD-L1 in a pulmonary *Mycobacterium avium* granuloma (Extended Data Fig. 6g). This supports that while PD-L1 expression could be a broader feature of certain granulomatous conditions, bright co-expression of IDO1 and PD-L1 may be specific to mycobacterial granulomas.

### Immunoregulatory features of granulomas are reflected in the peripheral blood of TB patients where they correlate to clinical progression and treatment status

The presence of immunosuppressive features in TB granulomas observed in our MIBI-TOF study has important implications for the treatment of TB infection. However, the invasive nature of procuring solid tissue limited the cohort size and scope by biasing towards advanced infections. Moreover, the single time point per patient in our tissue dataset precludes temporal analysis that would correlate disease severity with granuloma immunosuppressive features. Given the clinical feasibility of venous phlebotomy and large number of publicly available blood transcriptomic datasets, we sought to correlate these features in blood from TB patients. Therefore, we used MetaIntegrator to perform several multi-cohort analyses using peripheral blood transcriptome profiles from healthy subjects and patients with latent or active TB infection^59,60^.

We first determined if immune regulatory features identified in granulomas could be detected in blood by comparing publicly available gene expression data of patients with active TB (n = 647) to healthy controls (n = 197) from 13 independent cohorts (Fig. 6a). In line with PD-L1 and IDO1 expression in tissue data, significant and consistent upregulation of *IDO1* and *CD274* (PD-L1) was found in patients with active TB infection (effect size = 0.77 and 1.28, adj. p = 0.0009 and 0.006, respectively) (Fig. 6b). Additionally, checkpoint depletion in lymphoid cells was corroborated as well, with reduced expression of *PDCD1* (PD-1) and *LAG3* observed in the blood of active TB patients (effect size = −0.41 and −0.39, adj. p = 0.09 and 0.05 respectively).

**Figure 6.**
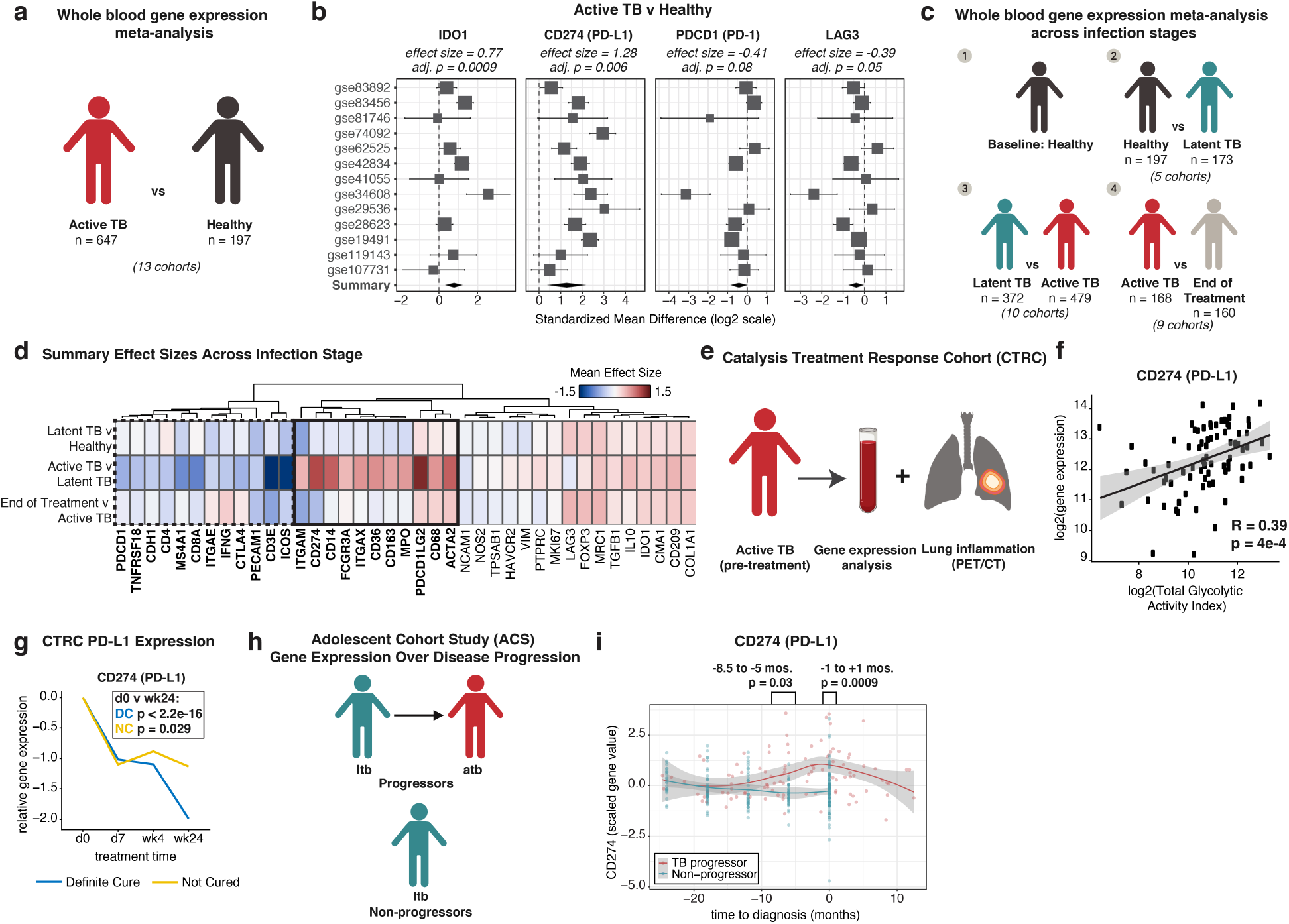
Immunoregulatory features of granulomas are reflected in the peripheral blood of TB patients where they correlate to clinical progression and treatment status. **(a)** Conceptual overview of the meta-analysis of patients with active TB (n = 647) versus healthy controls (n = 197). **(b)** Forest plots of gene expression differences in active TB versus healthy individuals. Cohort identifiers are shown on the y-axis. Boxes represent the standardized mean difference in gene expression (effect size). The size of the box is proportional to the sample size of that cohort. Whiskers represent the 95% confidence interval and diamonds (black) represent the overall difference in gene expression between two groups by integrating the standardized mean differences across all cohorts. The width of the diamond corresponds to its 95% confidence interval. The adj. p values (*q* values, FDR 5%) for the summary effect sizes are shown above each plot. **(c)** Conceptual overview of gene expression analysis across clinical infection stage. **(d)** Heatmap of summary gene expression (mean effect size) values in latent TB (n = 173) versus healthy controls (n = 197), latent TB (n = 372) versus active TB (n = 479), and active TB (n = 168) versus end-of-treatment (n = 160). Clinical stage is displayed on rows and genes are displayed across columns hierarchically clustered (Euclidean distance, complete linkage). Genes upregulated in active TB versus latent TB are shown in the solid black box, while downregulated genes are in the dashed black box. **(e)** Conceptual overview of the Catalysis Treatment Response Cohort (CTRC). **(f)** Correlation between PD-L1 gene expression and total glycolytic activity index (TGAI) represented as log2-transformed values. Linear fit (blue) with 95% confidence interval (grey) displayed. A Pearson correlation of 0.39 (p = 4 x 10^-4^ t-test) is displayed below the linear fit. **(g)** PD-L1 gene expression across treatment time broken down by cure status (blue = definite cure and yellow = no cure). Line represents mean expression in each time point, connected across time points. P-value determined with Student’s T-test for PD-L1 expression at d0 versus wk24 in the definite cure (DC, n = 71) and not-cured (NC, n = 7) groups. **(h)** Conceptual overview of the Adolescent Cohort Study (ACS). **(i)** PD-L1 gene expression in the ACS cohort across time prior to and after diagnosis of active TB stratified by progressors (red, n = 34) and non-progressors (blue, n = 109). Grey silhouette represents the 95% confidence interval. P-values determined by Welch two sample t-test.

Next, we analyzed transcriptomic data from 1,549 patients across 24 cohorts in order to determine if these features were specific to active infections. In line with the solid tissue analysis, differential expression of genes associated with regulatory myeloid cells (e.g. PDL1, PDL2, CD11b, CD11c, CD163) or T-cell immune checkpoint (e.g. PD1, CTLA4) delineated active from latent infections (Fig. 6d). Moreover, the majority of these genes returned to baseline levels seen in healthy controls after completing antimicrobial therapy (Fig. 6d, Extended Data Fig. 6h). Taken together, these results suggest a shift toward myeloid-mediated immune suppression that is specifically manifested during active TB infection.

Because PD-L1 gene expression exhibited the largest effect size relative to healthy controls, we next analyzed the Catalysis Treatment Response Cohort (CTRC) to determine if PD-L1 could be used as a biomarker for estimating disease burden and predicting clearance of infection (Fig 6e). Patients enrolled in this study provided venous blood and underwent PET/CT imaging^61,62^. Expression of PD-L1 at diagnosis was found to be directly correlated with total glycolytic activity index (TGAI), a radiographic metric for quantifying lung inflammation (Fig. 6f, Pearson r = 0.39 p = 4×10^-4^). 24 weeks after treatment, patients were clinically classified into four groups: definite cure, probable cure, possible cure, and not cured. Relative to the time of diagnosis, the reduction in PD-L1 expression in definitely cured patients (n = 71) were 2 times greater on average than in uncured patients (n = 7, Fig. 6g). A nearly identical trend was observed for PD-L2 as well (*PDCDLG2*, Extended Data Fig. 6i).

Lastly, we analyzed the Adolescent Cohort Study (ACS) to determine if PD-L1 could be used for predicting progression to active disease. Latently infected individuals enrolled in this study underwent regular blood collection and were clinically monitored for symptoms of active infection (Fig. 6h)^63,64^. PD-L1 transcript levels were significantly elevated in progressors (n = 34) relative to non-progressors (n = 109) 8.5 months prior to progression (p = 0.03) with the peak occurring at time of diagnosis (p = 0.0009) (Fig. 6i). Taken together, these results raise the intriguing possibility for using PD-L1 expression in peripheral blood as a predictive biomarker for early intervention in latently infected individuals.

## Discussion

After nearly 140 years of research into the pathophysiology of human TB infection, central questions remain unresolved, in large part because granuloma formation and progression are very difficult to emulate in tractable animal models. One of the most critical questions that remains unanswered is which immune mechanisms drive progression from latent infection to symptomatic active TB disease, the source of 11 million new cases and 1.5 million fatalities each year^1^. Furthermore, individual granuloma fate can vary significantly between lesions within individuals, raising questions about the local immune dynamics that influence a granuloma’s capacity to control infection while mitigating tissue damage. With this in mind we used MIBI-TOF to identify recurrent features of immune regulation in archival tissue from patients with active TB infection. This spatial cell atlas allowed us to relate granuloma structure and composition. We identified 19 unique cell subsets that preferentially organize into eight reoccurring cellular microenvironments. TB granulomas appear to follow a consistent structural outline characterized by spatially coordinated expression of PD-L1 and IDO1, myeloid core-infiltrating Tregs, and a striking absence of T cell activation as measured through PD-1 and Lag3. Some of these features, such as high expression of PD-L1 and presence of MDSC-like macrophages, were present in non-infectious granulomas as well pointing to certain universal immune programs associated with the granulomatous immune response. However, even compared to sarcoidosis, foreign body uptake, xanthomatosis, and endometrial lesions, spatially coordinated expression of IDO1 and PD-L1 was unique to mycobacterial granulomas.

Previous studies in the cynomolgus macaque model of TB have demonstrated that granulomas from a single individual can have disparate outcomes with respect to bacterial burden and inflammatory trajectory^10,11^. The variation we see in our imaging dataset suggests these local outcomes may be driven in part by unique cellular infiltrate and structure within each granuloma. We observed that certain features, such as a high frequency of CD8^+^ T cells, corresponded with reduced levels of more differentiated macrophage phenotypes, a profile consistently present in therapeutic resections where PD-L1 expression was also diminished. Since CD8^+^ T cells have been shown to be critical for clearance of TB infection^65,66^, understanding the immunological environments that promote CD8^+^ T cell activity could reveal novel insights into immune features critical for bacterial clearance.

By leveraging 29 publicly-available gene expression studies of over 1500 TB patients and healthy controls, we were able to identify these immunoregulatory signatures in peripheral blood and correlate them with disease burden and clinical outcome. Genes found in solid tissue to be overexpressed at the protein level by immunosuppressive myeloid cells (PD-L1, IDO1, CD163) were upregulated in blood. Similarly, genes associated with T-cell activation were downregulated, consistent with the rare incidence of PD-1 or Lag3 expression in tissue. Importantly, the magnitude of these trends was distinctly higher in patients with active disease relative to those with latent or treated infections. The highest effect size of genes measured in our analysis was observed for PD-L1. PD-L1 was found to correlate with pulmonary disease burden and appears to be a prognostic biomarker of progression from latent to active TB disease. Taken together, the work presented here reveals new aspects of immune regulation in TB infection that have important implications for understanding disease pathogenesis and improving clinical management.

The high levels of PD-L1 and IDO1 observed in the near absence of PD-1 offers clues into how the immunosuppressive niche during human infection is initiated and maintained. These findings are consistent with a TGFβ or IL-10 driven process where production of these cytokines can suppress inflammation, promote immunosuppression, and induce peripheral regulatory T cell differentiation and proliferation^67–70^. This is supported by recent work in mice where focal secretion of TGF-β within the myeloid core was suggested to preferentially suppress neighboring T cells and in non-human primates where granuloma formation was associated with IL-10 secretion ^71,72^.

Both IDO1 and PD-L1 have been shown to dampen anti-tumor immune responses in cancer, which has prompted the development of host-directed immunotherapies^73^. Our findings suggest that similar approaches could be used to block PD-L1 mediated immune suppression to promote bacterial clearance. However, evidence of T-cell activation or exhaustion is not present in our dataset or in the cynomolgus macaque model. This suggests that unlike checkpoint blockade in the setting of cancer, the efficacy of PD-L1 or PD-1 blockade could differ significantly. Recent reports of TB reactivation following PD-1 blockade illustrate the seemingly paradoxical effects that can occur with host directed therapies and emphasize the need to comprehensively map the temporal and spatial dynamics of these pathways^74–76^. In line with this, a critical next step will be to extend this work to relate these features to bacterial burden, inflammatory dynamics, and granuloma age in a primate model that accurately recapitulates human TB pathology.

To the best of our knowledge, this is the most comprehensive systems analysis of TB to date. We identified dynamics of cellular composition and immunoregulatory pathways in TB granulomas that are reflected in the peripheral immune response to TB. These results have implications both for developing host-directed immunotherapies and for identifying patients at risk of progression to active disease or treatment failure. Expression of proteins such as IDO1 and PD-L1 aligns with immune evasion mechanisms observed in tumor-immune microenvironment. The interface of granuloma and tumor immunobiology offers vast opportunities to explore how tactics of immune evasion in tumors may contribute to bacterial persistence in granulomas. Future multiplexed imaging studies of granulomas from controlled TB exposures will offer novel insights on how these local regulatory dynamics influence granuloma fate, and ultimately, infection outcome.

## Data and Code Availability

All custom code used to analyze data can be accessed at the following link: https://github.com/efrancis28/TBMIBI. All processed images and annotated single cell data will be made available on Mendeley Data.

## Methods

### Human Samples

Human samples were acquired in accordance with IRB protocol # 46646.

### Tuberculosis Granuloma Cohort

Formalin-fixed Paraffin-embedded (FFPE) *Mtb* infected tissues were acquired from Stanford Health Care’s tissue repository from seven patients undergoing a pre-treatment diagnostic biopsy. Tissues were screened to include those that were positive for Acid-Fast Bacillus (AFB+) and Mtb DNA by polymerase chain reaction (PCR). Archival surgical resection tissues were acquired from University of KwaZulu-Natal, Inkosi Albert Luthuli Central Hospital from six patients with *Mtb* infection who underwent therapeutic resection of infected tissue due to infection severity or treatment failure. This specimen group contained HIV^+^ patients on antiretroviral therapy (n = 3), HIV^-^ patients (n = 2), and one patient with no HIV infection reported. All clinical details for specimens can be found in Extended Data Table 1. 5 μm serial sections of each specimen were stained with hematoxylin and eosin (H<E) and inspected by an anatomic pathologist to screen for the presence of granulomatous inflammation. Regions with histologically solid granulomas or cellular granulomatous inflammation were included. Regions with excessive fibrosis or necrosis were excluded. Two 500 μm^2^ fields of view (FOV) were chosen from each tissue block for imaging.

### Non-tuberculous Granulomas and Controls Tissues

Regions of granulomatous inflammation from FFPE sarcoidosis and foreign body reactions from Stanford Health Care were chosen by an anatomic pathologist. 0.5 mm cores were selected and constructed into a tissue microarray (TMA). A pulmonary *Mycobacterium avium* case was acquired from Stanford Health Care through selection criteria of positive Acid-Fast Bacillus (AFB+) staining and PCR positivity for *M. avium Complex* (MAC). A 5 μm serial section of this specimen was stained with hematoxylin and eosin (H&E) and inspected by an anatomic pathologist to screen for the presence of granulomatous inflammation. Control tissues of FFPE tonsil, spleen, and placenta were acquired from Stanford Health Care. Small regions of each tissue were selected and placed in a TMA.

### Antibody Preparation

Antibodies were conjugated to isotopic metal reporters as described previously^19^. Following conjugation antibodies were diluted in Candor PBS Antibody Stabilization solution (Candor Bioscience). Antibodies were either stored at 4°C or lyophilized in 100 mM D-(+)-Trehalose dehydrate (Sigma Aldrich) with ultrapure distilled H_2_O for storage at −20°C. Prior to staining, lyophilized antibodies were reconstituted in a buffer of Tris (Thermo Fisher Scientific), sodium azide (Sigma Aldrich), ultrapure water (Thermo Fisher Scientific), and antibody stabilizer (Candor Bioscienc) to a concentration of 0.05 mg/mL. Information on the antibodies, metal reporters, and staining concentrations is located in Extended Data Table 1.

### Tissue Staining

Tissues were sectioned (5 μm section thickness) from tissue blocks on gold and tantalum-sputtered microscope slides. Slides were baked at 70 °C overnight followed by deparaffinization and rehydration with washes in xylene (3x), 100% ethanol (2x), 95% ethanol (2x), 80% ethanol (1x), 70% ethanol (1x), and ddH_2_O with a Leica ST4020 Linear Stainer (Leica Biosystems). Tissues next underwent antigen retrieval by submerging sides in 3-in-1 Target Retrieval Solution (pH 9, DAKO Agilent) and incubating at 97 °C for 40 minutes in a Lab Vision PT Module (Thermo Fisher Scientific). After cooling to room temperature slides were washed in 1x PBS IHC Washer Buffer with Tween 20 (Cell Marque) with 0.1% (w/v) bovine serum albumin (Thermo Fisher). Next, all tissues underwent two rounds of blocking, the first to block endogenous biotin and avidin with an Avidin/Biotin Blocking Kit (Biolegend). Tissues were then washed with wash buffer and blocked for 1 hour at room temperature with 1x TBS IHC Wash Buffer with Tween 20 with 3% (v/v) normal donkey serum (Sigma-Aldrich), 0.1% (v/v) cold fish skin gelatin (Sigma Aldrich), 0.1% (v/v) Triton X-100, and 0.05% (v/v) Sodium Azide. The first antibody cocktail was prepared in 1x TBS IHC Wash Buffer with Tween 20 with 3% (v/v) normal donkey serum (Sigma-Aldrich) and filtered through a 0.1 μm centrifugal filter (Millipore) prior to incubation with tissue overnight at 4 ° in a humidity chamber. Following the overnight incubation slides were washed twice for 5 minutes in wash buffer. The second day antibody cocktail was prepared as described and incubated with the tissues for 1 hour at 4 ° in a humidity chamber. Following staining, slides were washed twice for 5 minutes in wash buffer and fixed in a solution of 2% glutaraldehyde (Electron Microscopy Sciences) solution in low-barium PBS for 5 minutes. Slides were washed in PBS (1x), 0.1 M Tris at pH 8.5 (3x), ddH_2_O (2x), and then dehydrated by washing in 70% ethanol (1x), 80% ethanol (1x), 95% ethanol (2x), and 100% ethanol (2x). Slides were dried under vacuum prior to imaging.

### MIBI-TOF Imaging

Imaging was performed using a MIBI-TOF instrument with a Hyperion ion source. Xe^+^ primary ions were used to sequentially sputter pixels for a given FOV. The following imaging parameters were used:

- Acquisition setting: 80 kHz
- Field size: 500 μm^2^ (TB, *M. avium* and controls) or 450 μm^2^ (sarcoidosis) at 1024 x 1024 pixels
- Dwell time: 4 ms
- Median gun current on tissue: 1.45 nA Xe^+^
- Ion dose: 3.38 nAmp hours / mm^2^ for 500 μm^2^ FOVs and 3.75 nAmp hours / mm^2^ for 450 μm^2^

### Low-level Image Processing

Multiplexed image sets were extracted, slide background-subtracted, denoised, and aggregate filtered as previously described^19^. All parameters for these steps can be found in Extended Data Table 1. In addition to these processing steps, image compensation was performed to account for signal spillover due to adducts and oxides for the following interferences: Collagen-1 to IDO1 and Lag3, H3K9Ac to panCK and MPO, Chym/Tryp to MPO, Ki67 to CD209, CD20 to CD16, CD16 to IFNγ, CD11c to IDO1, and HLA-DR-DQ-DP to CD11b.

### Single Cell Segmentation

Nuclear segmentation was performed using an adapted version of the DeepCell^19–21^ CNN architecture. Training data was generated from MIBI-TOF images of human melanoma that were stained with HH3 to identify nuclei and Na^+^K^+^ATPase to identify the cell membrane. Color overlays of these two channels were used to manually segment nuclei in ImageJ. This generated training data with labels for the nuclear interior, nuclear border, and non-nuclear background. Training data was generated for 5 images to train the network architecture. Images were divided into overlapping crops of 64×64 pixels. Each crop was randomly flipped, rotated, and sheared during training to further augment the available data. Stochastic gradient descent was used to train the network, combined with early stopping to prevent over-fitting. Following training all MIBI-TOF images from our cohorts were provided as input to the network to predict the class of each pixel: nuclear interior, nuclear border, or non-nuclear background. The nuclear interior probability map for each image was thresholded and segmented as described previously^19^ followed by a 3-pixel radial expansion around each nucleus to define the cell object boundaries. A correction was applied to FOVs that contained multinucleated giant cells (MGNs). First each MGN was identified using a combination of HH3, CD45, and Vimentin and manually segmented in ImageJ to produce a binary mask of each MGN. All pixels within the binary mask were reassigned to belong to the MGN cell object(s).

### Single Cell Phenotyping and Composition

Single cell data was extracted for all cell objects and area-normalized. Cells with a sum of less than 0.1 area-normalized counts across all lineage channels were excluded from analysis. Single cell data was linearly scaled with a scaling factor of 100 and asinh-transformed with a co-factor of 5. All mass channels were scaled to 99.9^th^ percentile. In order to assign each cell to a lineage, the FlowSOM clustering algorithm was used in iterative rounds with the Bioconductor “FlowSOM” package in R^22^. The first clustering round separated cells into four major lineages using the “Metaclustering_consensus” function: immune, epithelial, fibroblast, and endothelial. Immune cells were then clustered again to delineate B cells, CD4^+^ T cells, CD8^+^ T cells, Tregs, neutrophils, mast cells, and mononuclear phagocytes (macrophages, monocytes, and dendritic cells). Immune cells with an expression profile not consistent with any of those subsets were annotated as ‘other immune.’ Lastly, the mononuclear phagocytes were clustered to 25 metaclusters which were merged into 7 groups. Giant cells were manually identified. γδ T cells were annotated as T cells with CD3 signal greater than or equal to the mean expression of CD4^+^ T cells and TCR-δ signal > 0.5 normalized expression. CD163 macrophages were identified as those with CD163 signal > 0.5 normalized expression. The relative abundance of all major lineages was determined out of total cells per FOV and the relative frequency of immune cell subsets was determined out of total immune cells per FOV.

### Immune Cell Composition Clustering and Cell Type Association Analysis

The Pearson correlation coefficient was calculated between all pairs of TB FOVs based on the relative frequency of all immune cell subsets. The coefficients were used to hierarchically cluster the FOVs using complete linkage and a distance metric of 1-correlation coefficient. To identify consensus clusters the percent variance explained was measured across a range of 1-10 clusters. The elbow point of this plot was used to determine the optimal number of clusters. A randomized dataset was produced to compare to the observed clustering by randomizing the frequency values across immune cell subsets within each FOV. This dataset was also clustered using the Pearson correlation coefficient and compared with the observed result. To assess the significance of co-occurrence of cell types, a chi-square test was run between all cell type pairs using the counts of each cell type across all TB FOVs. The resulting p-values were adjusted using a false discovery rate (FDR) of 5%.

### Protein Enrichment Analysis

A spatial enrichment approached was used as previously described^19^ to identify patterns of protein enrichment or exclusion across all protein pairs. HH3, Na^+^K^+^ATPase, and HLA Class 1 were excluded from the analysis. For each pair of markers, X and Y, the number of times cells positive for protein X was within a ~50 um radius of cells positive for protein Y was counted. Thresholds for positivity were customized to each marker individually using a silhouette scanning approach from the MetaCyto software in R^77^. A null distribution was produced by performing 1000 bootstrap permutations where the locations of cells positive for protein Y were randomized. A z-score was calculated comparing the number of true cooccurrences of cells positive for protein X and Y relative to the null distribution. For each pair of proteins X and Y the average z-score was calculated across all TB FOVs. Next, all positive enrichments between protein pairs (average Z score > 0, excluded self-self protein enrichment scores) were used to produce a weighted, undirected network where the nodes are the individual markers and the edge weights are proportional to the average z-score (higher z-score à shorter edge length). The leading eigenvector algorithm for community detection was used to identify protein enrichment communities in this network^78^.

### Spatial Latent Dirichlet Allocation

Spatial-LDA is an adaptation of Latent Dirichlet Allocation (LDA), a machine learning approach used to model topics in text documents, where topics consist of words with a high probability of cooccurrence, allowing mapping of topics to abstract definitions (ex. [‘dog’, ‘frog’, ‘horse] à ‘animals’). Spatial-LDA builds on this paradigm by representing CPMs as documents and cell types as words. Spatial-LDA was conducted to identify topics (here referred to as microenvironments) across all TB FOVs. Cell types with fewer than 100 cells across the entire cohort were excluded from analysis (γδ T cells and CD209^+^ DC). Furthermore, multinucleated giant cells were excluded due to their cell size. Spatial-LDA was implemented as described^30^ with d = 1000, a spatial radius r = 50 μm to complement the protein enrichment analysis, and a microenvironment (ME) number of n = 8. The ME number was determined empirically. For each FOV a maximum probability map (MaxPM) was produced by classifying each cell to the microenvironment with the highest probability and coloring that cell by its microenvironment and probability. The cell type preferences for each ME were determined by assessing the mean probability for all cell types across all MEs. The mean expression for each functional marker across MEs was determined by weighting protein expression by ME probability and calculating the mean of weighted expression values across markers and MEs. The mean probability for all MEs was determined for all FOVs (average of single cell values) and used to conduct a Principal Component Analysis (PCA). The clustering approach described for immune cell frequency clusters (above) was applied to ME frequencies across FOVs to identify the optimal number of ME clusters to capture the maximal variance in our dataset.

### Identification of the Myeloid Core

In order to assess which microenvironments represented the histologically defined myeloid core, binary masks of the myeloid core were generated for 18/26 FOVs. The masks were generated by first combining the signal of CD11c, CD11b, CD14, CD206, CD68, and PD-L1. The combined images were imported into ImageJ and hand-annotated using ROI annotation tools. The annotated ROI was exported as a binary mask. This mask was further processed in Matlab to close any holes, exclude objects smaller than 1000 pixels, and dilate the mask to smooth edges. Next cells were assigned to belonging to the myeloid core if they had complete overlap with the binary mask. Cells on the mask boundary or outside of the mask were designated as ‘non-myeloid core.’ The proportion of cells in the myeloid core was assessed across each ME for the 18 FOVs and a MEs with a median frequency in the myeloid core > 50% were designated as myeloid core MEs.

### Myeloid Cell UMAP Visualization

UMAP embeddings were determined for all non-granulocytic myeloid cells using the R implementation^79^ with the following parameters: n_neighbors = 15, min_dist = 0.1 and the following markers: CD45, CD68, CD206, CD11c, CD11b, CD14, CD16, CD209, and CD163.

### Immunoregulatory Protein Analysis

Positivity thresholds for IDO1, PD-L1, PD-1, and Lag3 were automatically determined as described above. Immune control tissues tonsil, spleen, and placenta were used to validate antibody performance. Correlation between IDO1 and PD-L1 was determined across the entire cohort and subsets of specimens using Pearson correlation analysis. The frequency of cells positive for IDO1 and PD-L1 were enumerated across all subsets. To assess PD-L1 and IDO1 positivity with respect to ME and cell subset, the total number of cells across all myeloid subsets per ME was pooled across all FOVs. The quantity of cells for each subset positive for IDO1 or PD-L1 was determined per ME. Any ME with < 1% of the total for a subset was excluded from analysis. PD-1 and Lag3 expression were analyzed on lymphocytes or total immune cells. PD-1 and Lag3 were also analyzed on immune cells from a human Triple Negative Breast Cancer (TNBC) cohort that was previously published by our group^19^. Positivity for PD-1 and Lag3 for TNBC immune cells was determined as described in the originally published study.

### Cell Composition Analysis of Sarcoidosis and Tuberculosis

Single cells from sarcoidosis FOVs were segmented as described above. Single cell data was extracted, transformed, and normalized along with TB single cell data. Single cells were included in the described FlowSOM clustering procedure. To compare the cellular diversity of TB with sarcoidosis the Shannon Diversity Index was calculated using the counts of all cell subsets per TB or sarcoidosis FOV.

### Immunohistochemistry of PD-L1 and IDO1

Immunohistochemistry (IHC) for PD-L1 and IDO1 was performed using the antibody reagents in Extended Data Table 1 at a concentration of 1 μg/mL. The IHC protocol mirrors the MIBI-TOF protocol with the addition of blocking endogenous peroxidase activity with 3% H_2_O_2_ (Sigma Aldrich) in ddH_2_O after epitope retrieval. On the second day of staining, instead of proceeding with the MIBI-TOF protocol, tissues were washed twice for 5 minutes in wash buffer and stained using the ImmPRESS universal (Anti-Mouse/Anti-Rabbit) kit (Vector labs).

### Whole Blood Transcriptomic Analysis

Publicly available gene expression data sets (Extended Data Table 2) were collected, annotated, and used for meta-analysis conducted using MetaIntegrator^59^. Gene expression matrices were prepared for each dataset to determine effect sizes for genes of all proteins included in the MIBI-TOF analysis and an additional set of genes with similar biological function, such as *ICOS* and *CTLA4*. Summary effect sizes were calculated to assess gene expression differences across clinical groups (healthy, active TB, latent TB, end of treatment, TB progression, and during treatment). For the Catalysis Treatment Response Cohort (CTRC) gene expression measurements at diagnosis of TB were correlated with matched Total Glycolytic Activity Index (TGAI), a readout of PET-CT activity. A linear regression was fit between *CD274* gene expression and *TGAI* and the correlation was assessed with Pearson correlation analysis. To assess *CD274* and *PDCDLG2* gene expression over treatment, expression values were normalized to the measurement taken at diagnosis (d0). Gene expression data in the Adolescent Cohort Study (ACS) were separated by progression status. Local regression (LOESS) was used to fit the gene expression data over time in each group. The significance of separation between progressors and non-progressors was determined in two different time intervals using a Student’s t-test.

### Software

Image processing was conducted with Matlab 2016a and Matlab 2019b. Statistical analysis was conducted in Matlab 2016a, Matlab 2019b, and R. Data visualization and plots were generated in R. Representative images were processed in Adobe Photoshop and figures were prepared in Adobe Illustrator. Schematic visualizations were produced with Biorender.

## Author Contributions

E.F.M conceived the study design, performed experiments, analyzed data, and wrote the manuscript. M.D. conducted the blood transcriptomics analysis. L.K. assisted with analysis conceptualization. Z.C. and V.J. implemented the spatial-LDA analysis. L.K, A.D., N.F.G., A.B., and W.G. assisted with data analysis. M.B. assisted with assay development. D.V.V. developed Deepcell. M.F., P.K.R., E.F., M.V.D.R., N.B., and A.J.C.S. provided samples and consulted on tissue cohort design. S.C.B, P.K., and M.A. supervised the work.

## Acknowledgments

The authors thank Tyler Risom, David Glass, Matthew Carter, and Anne Kasmar for discussions and comments. The authors thank Pauline Chu and the Stanford Human Histology Core for providing technical assistance. E.F.M was supported by the NSF Graduate Research Fellowship (grant no. 2017242837). L.K. was a Damon Runyon Fellow supported by the Damon Runyon Cancer Research Foundation (DRG-2292-17) and a non-stipendiary awardee of the EMBO Long-Term fellowship (ALTF 1128–2016). Noah Greenwald was supported by F31CA246880. A.J.C.S. was supported R61/33AI138280, R01AI134810, the CRDF Global, the South African Medical Research Council, and an NRF BRICS Multilateral grant to A.J.C.S. M.A. was supported by 1-DP5-OD019822. S.C.B. and M.A. were jointly supported by 1R01AG056287 and 1R01AG057915, 1U24CA224309, the Bill and Melinda Gates Foundation, and a Translational Research Award from the Stanford Cancer Institute. S.J.G. was supported by U19 AI104209, R01 AR067145, and R01 AI32494.

## References

1. WHO | Global tuberculosis report 2019. WHO (2020).

2. Cohen, S. B. et al. Alveolar Macrophages Provide an Early Mycobacterium tuberculosis Niche and Initiate Dissemination. Cell Host Microbe 24, 439–446.e4 (2018).

3. Wolf, A. J. et al. Mycobacterium tuberculosis Infects Dendritic Cells with High Frequency and Impairs Their Function In Vivo. J. Immunol. 179, 2509–2519 (2007).

4. Bold, T. D. & Ernst, J. D. Who benefits from granulomas, mycobacteria or host? Cell 136, 17–9 (2009).

5. Davis, J. M. & Ramakrishnan, L. The Role of the Granuloma in Expansion and Dissemination of Early Tuberculous Infection. Cell 136, 37–49 (2009).

6. Ramakrishnan, L. Revisiting the role of the granuloma in tuberculosis. Nat. Rev. Immunol. 12, (2012).

7. Cadena, A. M., Fortune, S. M. & Flynn, J. L. Heterogeneity in tuberculosis. (2017) doi:10.1038/nri.2017.69.

8. Subbian, S. et al. Lesion-Specific Immune Response in Granulomas of Patients with Pulmonary Tuberculosis: A Pilot Study. PLoS One 10, e0132249 (2015).

9. Coleman, M. T. et al. Early Changes by (18)Fluorodeoxyglucose positron emission tomography coregistered with computed tomography predict outcome after Mycobacterium tuberculosis infection in cynomolgus macaques. Infect. Immun. 82, 2400–4 (2014).

10. Lin, P. L. et al. Sterilization of granulomas is common in active and latent tuberculosis despite within-host variability in bacterial killing. Nat. Med. 20, 75–79 (2013).

11. Martin, C. J. et al. Digitally Barcoding Mycobacterium tuberculosis Reveals In Vivo Infection Dynamics in the Macaque Model of Tuberculosis. MBio 8, e00312–17 (2017).

12. Carow, B. et al. Spatial and temporal localization of immune transcripts defines hallmarks and diversity in the tuberculosis granuloma. Nat. Commun. 10, 1–15 (2019).

13. Marakalala, M. J. et al. Inflammatory signaling in human tuberculosis granulomas is spatially organized. Nat. Med. 22, 531–538 (2016).

14. Wong, E. A. et al. Low levels of T cell exhaustion in tuberculous lung granulomas. Infect. Immun. 86, (2018).

15. Kauffman, K. D. et al. Defective positioning in granulomas but not lung-homing limits CD4 T-cell interactions with Mycobacterium tuberculosis-infected macrophages in rhesus macaques. Mucosal Immunol. 11, 462–473 (2018).

16. Keren, L. et al. MIBI-TOF: A multiplexed imaging platform relates cellular phenotypes and tissue structure. Sci. Adv. 5, eaax5851 (2019).

17. Krishnan, N., Robertson, B. D. & Thwaites, G. The mechanisms and consequences of the extra-pulmonary dissemination of Mycobacterium tuberculosis. Tuberculosis vol. 90 361–366 (2010).

18. Ranaivomanana, P. et al. Cytokine Biomarkers Associated with Human Extra-Pulmonary Tuberculosis Clinical Strains and Symptoms. Front. Microbiol. 9, 275 (2018).

19. Keren, L. et al. A Structured Tumor-Immune Microenvironment in Triple Negative Breast Cancer Revealed by Multiplexed Ion Beam Imaging. Cell 174, 1373–1387.e19 (2018).

20. Van Valen, D. A. et al. Deep Learning Automates the Quantitative Analysis of Individual Cells in Live-Cell Imaging Experiments. PLOS Comput. Biol. 12, e1005177 (2016).

21. Bannon, D. et al. Dynamic allocation of computational resources for deep learning-enabled cellular image analysis with Kubernetes. bioRxiv 505032 (2019) doi:10.1101/505032.

22. Van Gassen, S. et al. FlowSOM: Using self-organizing maps for visualization and interpretation of cytometry data. Cytom. Part A 87, 636–645 (2015).

23. Flynn, J. L., Chan, J. & Lin, P. L. Macrophages and control of granulomatous inflammation in tuberculosis. Mucosal Immunology vol. 4 271–278 (2011).

24. Garcia-Rodriguez, K. M., Goenka, A., Alonso-Rasgado, M. T., Hernández-Pando, R. & Bulfone-Paus, S. The role of mast cells in tuberculosis: Orchestrating innate immune crosstalk? Frontiers in Immunology vol. 8 (2017).

25. Ratnam, S., Ratnam, S., Puri, B. K. & Chandrasekhar, S. Mast cell response during the early phase of tuberculosis: an electron microscopic study. Can. J. Microbiol. 23, 1245–1251 (1977).

26. Carlos, D. et al. Mast Cells Modulate Pulmonary Acute Inflammation and Host Defense in a Murine Model of Tuberculosis. J. Infect. Dis. 196, 1361–1368 (2007).

27. Polena, H. et al. Mycobacterium tuberculosis exploits the formation of new blood vessels for its dissemination. Sci. Rep. 6, 1–11 (2016).

28. Oehlers, S. H. et al. Interception of host angiogenic signalling limits mycobacterial growth. Nature 517, 612–615 (2015).

29. Girvan, M. & Newman, M. E. J. Community structure in social and biological networks. Proc. Natl. Acad. Sci. U. S. A. 99, 7821–7826 (2002).

30. Chen, Z., Soifer, I., Hilton, H., Keren, L. & Jojic, V. Modeling Multiplexed Images with Spatial-LDA Reveals Novel Tissue Microenvironments. J. Comput. Biol. (2020) doi:10.1089/cmb.2019.0340.

31. Shi, J. et al. PD-1 Controls Follicular T Helper Cell Positioning and Function. Immunity 49, 264–274.e4 (2018).

32. Dieu-Nosjean, M. C., Goc, J., Giraldo, N. A., Sautès-Fridman, C. & Fridman, W. H. Tertiary lymphoid structures in cancer and beyond. Trends in Immunology vol. 35 571–580 (2014).

33. Ulrichs, T. et al. Human tuberculous granulomas induce peripheral lymphoid follicle-like structures to orchestrate local host defence in the lung. J. Pathol. 204, 217–228 (2004).

34. Difazio, R. M. et al. Active transforming growth factor-β is associated with phenotypic changes in granulomas after drug treatment in pulmonary tuberculosis. DARU, J. Pharm. Sci. 24, 6 (2016).

35. Krystel-Whittemore, M., Dileepan, K. N. & Wood, J. G. Mast cell: A multi-functional master cell. Frontiers in Immunology vol. 6 (2016).

36. Shen, L. et al. PD-1/PD-L pathway inhibits M.tb-specific CD4+ T-cell functions and phagocytosis of macrophages in active tuberculosis. Sci. Rep. 6, 38362 (2016).

37. Shen, L. et al. The characteristic profiles of PD-1 and PD-L1 expressions and dynamic changes during treatment in active tuberculosis. Tuberculosis 101, 146–150 (2016).

38. Mehra, S. et al. Granuloma correlates of protection against tuberculosis and mechanisms of immune modulation by Mycobacterium tuberculosis. J. Infect. Dis. 207, 1115–27 (2013).

39. Munn, D. H. & Mellor, A. L. IDO in the Tumor Microenvironment: Inflammation, Counter-Regulation, and Tolerance. Trends in Immunology vol. 37 193–207 (2016).

40. Gautam, U. S. et al. In vivo inhibition of tryptophan catabolism reorganizes the tuberculoma and augments immune-mediated control of Mycobacterium tuberculosis. Proc. Natl. Acad. Sci. U. S. A. 115, E62–E71 (2018).

41. Jurado, J. O. et al. Programmed death (PD)-1 :PD-ligand 1/PD-ligand 2 pathway inhibits T cell effector functions during human tuberculosis. J. Immunol. 181, 116–25 (2008).

42. Human myeloid derived suppressor cell (MDSC) subset phenotypes I The Journal of Immunology. https://www.jimmunol.org/content/198/1_Supplement/211.2.

43. Gideon, H. P., Phuah, J., Junecko, B. A. & Mattila, J. T. Neutrophils express pro-and anti-inflammatory cytokines in granulomas from Mycobacterium tuberculosis-infected cynomolgus macaques. Mucosal Immunol. 12, 1370–1381 (2019).

44. Kanamori, M., Nakatsukasa, H., Okada, M., Lu, Q. & Yoshimura, A. Induced Regulatory T Cells: Their Development, Stability, and Applications. Trends in Immunology vol. 37 803–811 (2016).

45. Hsu, P. et al. IL-10 Potentiates Differentiation of Human Induced Regulatory T Cells via STAT3 and Foxo1. J. Immunol. 195, 3665–3674 (2015).

46. Llopiz, D. et al. IL-10 expression defines an immunosuppressive dendritic cell population induced by antitumor therapeutic vaccination. Oncotarget 8, 2659–2671 (2017).

47. Ribeiro-Rodrigues, R. et al. A role for CD4+CD25+ T cells in regulation of the immune response during human tuberculosis. Clin. Exp. Immunol. 144, 25–34 (2006).

48. Scott-Browne, J. P. et al. Expansion and function of Foxp3-expressing T regulatory cells during tuberculosis. J. Exp. Med. 204, 2159–2169 (2007).

49. Guyot-Revol, V., Innes, J. A., Hackforth, S., Hinks, T. & Lalvani, A. Regulatory T cells are expanded in blood and disease sites in patients with tuberculosis. Am. J. Respir. Crit. Care Med. 173, 803–810 (2006).

50. Green, A. M. et al. CD4 + Regulatory T Cells in a Cynomolgus Macaque Model of Mycobacterium tuberculosis Infection. J. Infect. Dis. 202, 533–541 (2010).

51. Bagaitkar, J. Cellular dynamics of resolving inflammation. Blood vol. 124 1701–1703 (2014).

52. Wherry, E. J. & Kurachi, M. Molecular and cellular insights into T cell exhaustion. Nature Reviews Immunology vol. 15 486–499 (2015).

53. Baughman, R. P., Lower, E. E. & Du Bois, R. M. Sarcoidosis. in Lancet vol. 361 1111–1118 (Elsevier Limited, 2003).

54. Koth, L. L. et al. Sarcoidosis blood transcriptome reflects lung inflammation and overlaps with tuberculosis. Am. J. Respir. Crit. Care Med. 184, 1153–1163 (2011).

55. Maertzdorf, J. et al. Common patterns and disease-related signatures in tuberculosis and sarcoidosis. Proc. Natl. Acad. Sci. U. S. A. 109, 7853–7858 (2012).

56. Wahlström, J. et al. Analysis of intracellular cytokines in CD4+ and CD8+ lung and blood T cells in sarcoidosis. Am. J. Respir. Crit. Care Med. 163, 115–121 (2001).

57. Rossi, G. A. et al. Helper T-lymphocytes in pulmonary sarcoidosis. Functional analysis of a lung T-cell subpopulation in patients with active disease. Am. Rev. Respir. Dis. 133, 1086–1090 (1986).

58. Facco, M. et al. Sarcoidosis is a Th1/Th17 multisystem disorder. Thorax 66, 144–150 (2011).

59. Haynes, W. A. et al. Empowering multi-cohort gene expression analysis to increase reproducibility. in Pacific Symposium on Biocomputing vol. 0 144–153 (World Scientific Publishing Co. Pte Ltd, 2017).

60. Sweeney, T. E., Haynes, W. A., Vallania, F., Ioannidis, J. P. & Khatri, P. Methods to increase reproducibility in differential gene expression via meta-analysis. Nucleic Acids Res. (2017) doi:10.1093/nar/gkw797.

61. Roy Chowdhury, R. et al. A multi-cohort study of the immune factors associated with M. tuberculosis infection outcomes. Nature vol. 560 644–648 (2018).

62. Malherbe, S. T. et al. Persisting positron emission tomography lesion activity and Mycobacterium tuberculosis mRNA after tuberculosis cure. Nat. Med. 22, 1094–1100 (2016).

63. Scriba, T. J. et al. Sequential inflammatory processes define human progression from M. tuberculosis infection to tuberculosis disease. PLOS Pathog. 13, e1006687 (2017).

64. Zak, D. E. et al. A blood RNA signature for tuberculosis disease risk: a prospective cohort study. Lancet 387, 2312–2322 (2016).

65. Flynn, J. L., Goldstein, M. M., Triebold, K. J., Koller, B. & Bloom, B. R. Major histocompatibility complex class I-restricted T cells are required for resistance to Mycobacterium tuberculosis infection. Proc. Natl. Acad. Sci. U. S. A. 89, 12013–12017 (1992).

66. Lalvani, A. et al. Human cytolytic and interferon γ-secreting CD8+ T lymphocytes specific for Mycobacterium tuberculosis. Proc. Natl. Acad. Sci. U. S. A. 95, 270–275 (1998).

67. Salazar-Onfray, F., López, M. N. & Mendoza-Naranjo, A. Paradoxical effects of cytokines in tumor immune surveillance and tumor immune escape. Cytokine Growth Factor Rev. 18, 171–182 (2007).

68. Li, M. O., Sanjabi, S. & Flavell, R. A. A. Transforming Growth Factor-β Controls Development, Homeostasis, and Tolerance of T Cells by Regulatory T Cell-Dependent and -Independent Mechanisms. Immunity 25, 455–471 (2006).

69. Jarnicki, A. G., Lysaght, J., Todryk, S. & Mills, K. H. G. Suppression of Antitumor Immunity by IL-10 and TGF-β-Producing T Cells Infiltrating the Growing Tumor: Influence of Tumor Environment on the Induction of CD4 + and CD8 + Regulatory T Cells. J. Immunol. 177, 896–904 (2006).

70. Letterio, J. J. & Roberts, A. B. REGULATION OF IMMUNE RESPONSES BY TGF-β. Annu. Rev. Immunol. 16, 137–161 (1998).

71. Gern, B. H. et al. TGFβ restricts T cell function and bacterial control within the tuberculous granuloma. bioRxiv 696534 (2019) doi:10.1101/696534.

72. Wong, E. A. et al. IL-10 Impairs Local Immune Response in Lung Granulomas and Lymph Nodes during Early Mycobacterium tuberculosis Infection. J. Immunol. 204, 644–659 (2020).

73. Havel, J. J., Chowell, D. & Chan, T. A. The evolving landscape of biomarkers for checkpoint inhibitor immunotherapy. Nature Reviews Cancer vol. 19 133–150 (2019).

74. Barber, D. L. et al. Tuberculosis following PD-1 blockade for cancer immunotherapy. Sci. Transl. Med. 11, (2019).

75. Anastasopoulou, A., Ziogas, D. C., Samarkos, M., Kirkwood, J. M. & Gogas, H. Reactivation of tuberculosis in cancer patients following administration of immune checkpoint inhibitors: Current evidence and clinical practice recommendations. Journal for ImmunoTherapy of Cancer vol. 7 239 (2019).

76. Tezera, L. B. et al. Anti-PD-1 immunotherapy leads to tuberculosis reactivation via dysregulation of TNF-α. Elife 9, (2020).

77. Hu, Z. et al. MetaCyto: A Tool for Automated Meta-analysis of Mass and Flow Cytometry Data. Cell Rep. 24, 1377–1388 (2018).

78. Newman, M. E. J. Finding community structure in networks using the eigenvectors of matrices. Phys. Rev. E - Stat. Nonlinear, Soft Matter Phys. 74, (2006).

79. McInnes, L., Healy, J. & Melville, J. UMAP: Uniform Manifold Approximation and Projection for Dimension Reduction. (2018).

