## Extended Data Figures for "Multiplexed imaging of human tuberculosis granulomas uncovers immunoregulatory features conserved across tissue and blood"

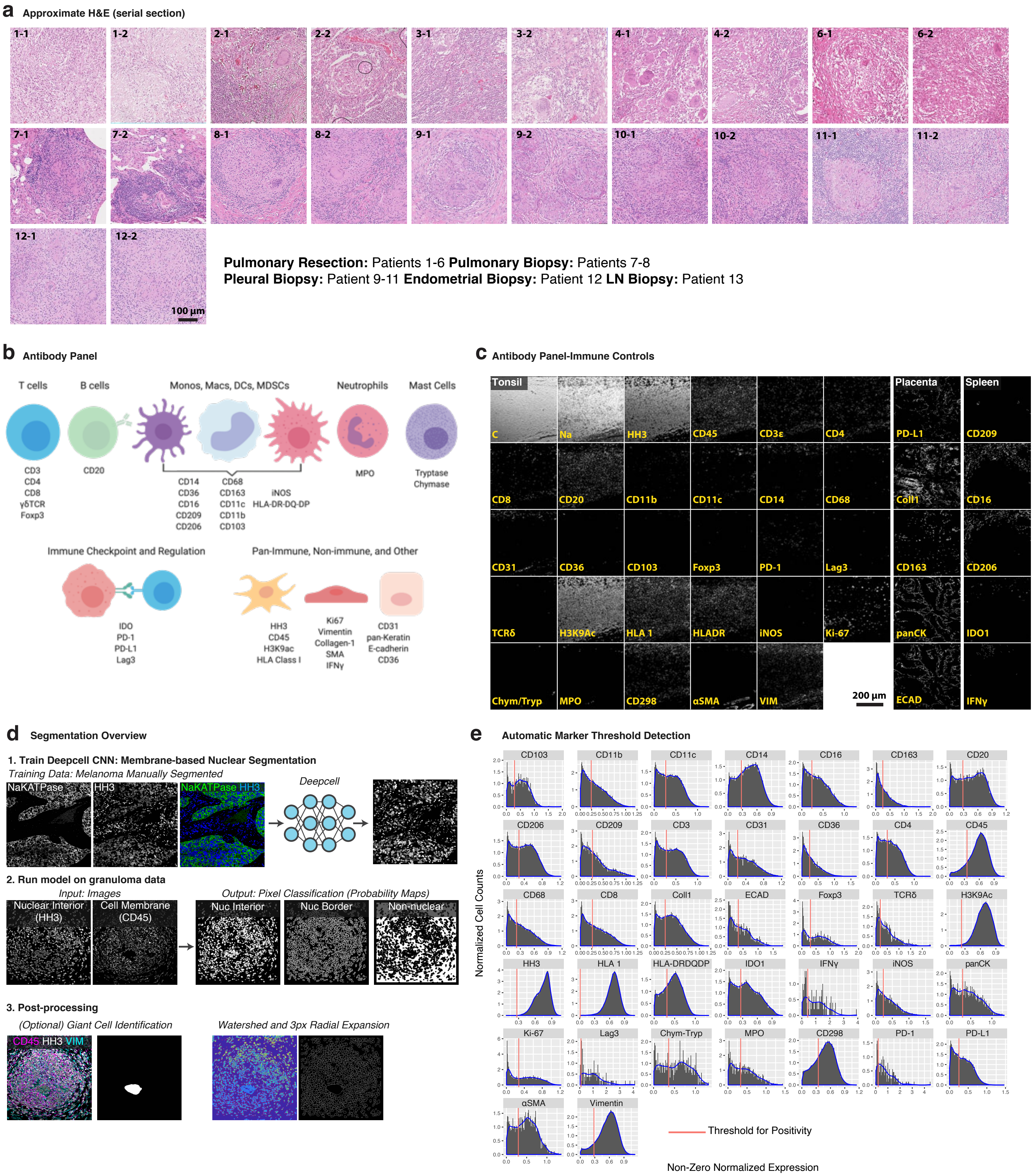

a Approximate H&E (serial section)
1-1
1-2
2-1
2-2
3-1
3-2
4-1
4-2
6-1
6-2
7-1
7-2
8-1
8-2
9-1
9-2
10-1
10-2
11-1
11-2
12-1
12-2
100 µm
**Pulmonary Resection:** Patients 1-6 **Pulmonary Biopsy:** Patients 7-8
**Pleural Biopsy:** Patient 9-11 **Endometrial Biopsy:** Patient 12 **LN Biopsy:** Patient 13
b Antibody Panel
T cells
B cells
Monos, Macs, DCs, MDSCs
Neutrophils
Mast Cells
CD3
CD4
CD8
γδTCR
Foxp3
CD20
CD14
CD36
CD16
CD209
CD206
CD68
CD163
CD11c
CD11b
CD103
iNOS
HLA-DR-DQ-DP
MPO
Tryptase
Chymase
Immune Checkpoint and Regulation
IDO
PD-1
PD-L1
Lag3
Pan-Immune, Non-immune, and Other
HH3
CD45
H3K9ac
HLA Class I
Ki67
Vimentin
Collagen-1
SMA
IFNγ
CD31
pan-Keratin
E-cadherin
CD36
c Antibody Panel-Immune Controls
Tonsil
Placenta
Spleen
C
Na
HH3
CD45
CD3ε
CD4
PD-L1
CD209
CD8
CD20
CD11b
CD11c
CD14
CD68
Coll1
CD16
CD31
CD36
CD103
Foxp3
PD-1
Lag3
CD163
CD206
TCRδ
H3K9Ac
HLA 1
HLADR
iNOS
Ki-67
panCK
IDO1
Chym/Tryp
MPO
CD298
αSMA
VIM
200 µm
ECAD
IFNγ
d Segmentation Overview
1. Train Deepcell CNN: Membrane-based Nuclear Segmentation
Training Data: Melanoma Manually Segmented
NaKATPase
HH3
NaKATPase HH3
Deepcell
2. Run model on granuloma data
Input: Images
Output: Pixel Classification (Probability Maps)
Nuclear Interior (HH3)
Cell Membrane (CD45)
Nuc Interior
Nuc Border
Non-nuclear
3. Post-processing
(Optional) Giant Cell Identification
Watershed and 3px Radial Expansion
CD45 HH3 VIM
e Automatic Marker Threshold Detection
CD103
CD11b
CD11c
CD14
CD16
CD163
CD20
CD206
CD209
CD3
CD31
CD36
CD4
CD45
CD68
CD8
Coll1
ECAD
Foxp3
TCRδ
H3K9Ac
HH3
HLA 1
HLA-DRDQDP
IDO1
IFNγ
iNOS
panCK
Ki-67
Lag3
Chym-Tryp
MPO
CD298
PD-1
PD-L1
αSMA
Vimentin
Normalized Cell Counts
Threshold for Positivity
Non-Zero Normalized Expression

**Extended Data Figure 1. Multiplexed imaging of human tuberculosis granulomas. (a)** Hematoxylin and eosin stained serial sections of FOVs for MIBI-TOF imaging. **(b)** Multiplexed antibody panel grouped by marker category. **(c)** Grayscale images of endogenous ion signal and proteins in control tissues (tonsil, spleen, placenta). **(d)** Workflow for Deepcell-based segmentation of single cells from multiplexed images. **(e)** Histograms of non-zero signal for all proteins from single cell data. Blue line represents Gaussian smoothed density fit of histogram. Red line represents automatically identified threshold for marker positivity.

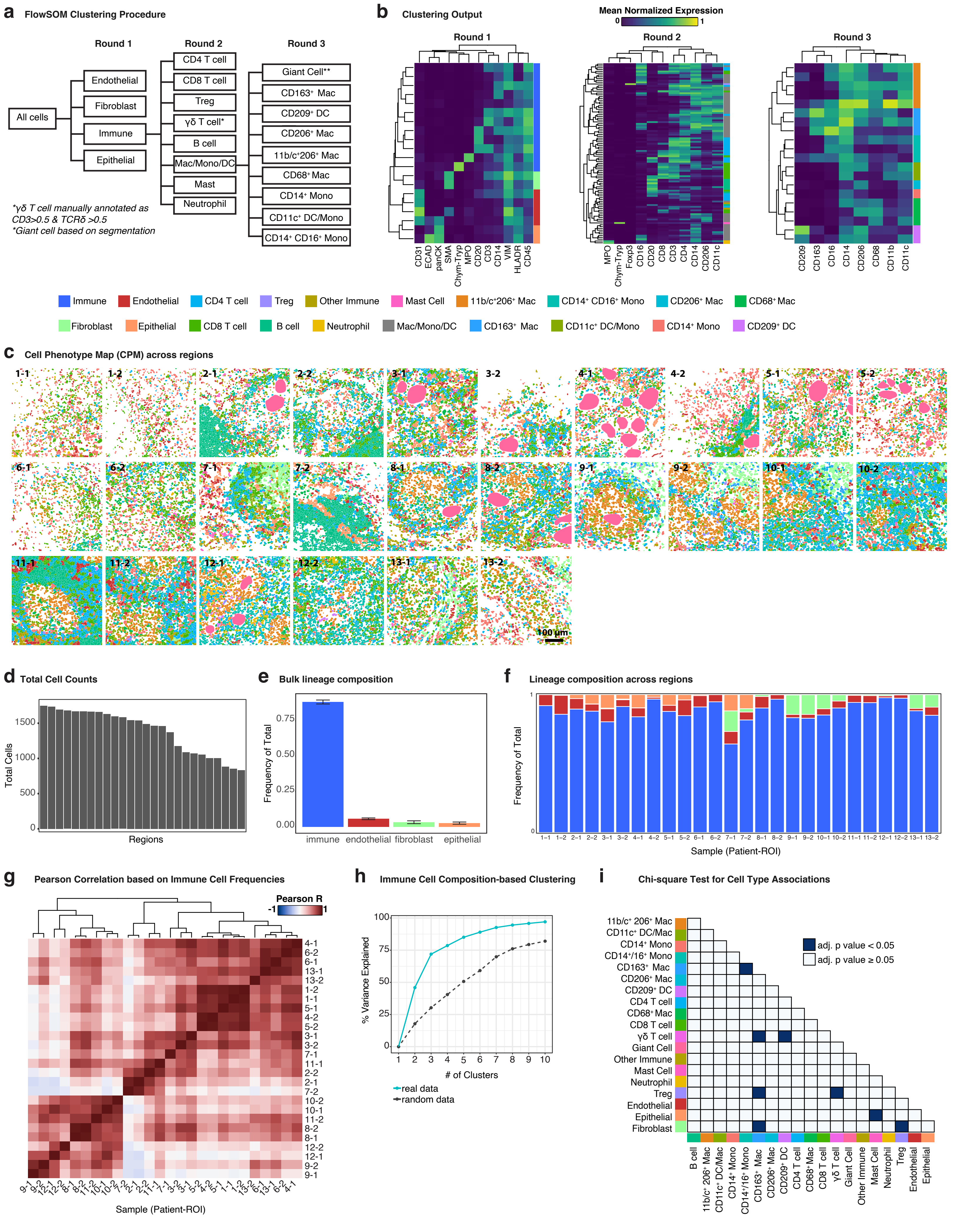
a
FlowSOM Clustering Procedure
Round 1
Round 2
Round 3
All cells
Endothelial
Fibroblast
Immune
Epithelial
CD4 T cell
CD8 T cell
Treg
γδ T cell*
B cell
Mac/Mono/DC
Mast
Neutrophil
Giant Cell**
CD163⁺ Mac
CD209⁺ DC
CD206⁺ Mac
11b/c⁺206⁺ Mac
CD68⁺ Mac
CD14⁺ Mono
CD11c⁺ DC/Mono
CD14⁺ CD16⁺ Mono
*γδ T cell manually annotated as CD3>0.5 & TCRδ >0.5
*Giant cell based on segmentation
b
Clustering Output
Mean Normalized Expression
0
1
Round 1
Round 2
Round 3
CD31
ECAD
panCK
SMA
Chym-Tryp
MPO
CD20
CD3
CD14
VIM
HLADR
CD45
MPO
Chym-Tryp
Foxp3
CD16
CD20
CD8
CD3
CD4
CD14
CD206
CD11c
CD209
CD163
CD16
CD14
CD206
CD68
CD11b
CD11c
Immune
Endothelial
CD4 T cell
Treg
Other Immune
Mast Cell
11b/c⁺206⁺ Mac
CD14⁺ CD16⁺ Mono
CD206⁺ Mac
CD68⁺ Mac
Fibroblast
Epithelial
CD8 T cell
B cell
Neutrophil
Mac/Mono/DC
CD163⁺ Mac
CD11c⁺ DC/Mono
CD14⁺ Mono
CD209⁺ DC
c
Cell Phenotype Map (CPM) across regions
1-1
1-2
2-1
2-2
3-1
3-2
4-1
4-2
5-1
5-2
6-1
6-2
7-1
7-2
8-1
8-2
9-1
9-2
10-1
10-2
11-1
11-2
12-1
12-2
13-1
13-2
100 µm
d
Total Cell Counts
Total Cells
1500
1000
500
0
Regions
e
Bulk lineage composition
Frequency of Total
0.75
0.50
0.25
0.00
immune
endothelial
fibroblast
epithelial
f
Lineage composition across regions
Frequency of Total
1
0
1–1 1–2 2–1 2–2 3–1 3–2 4–1 4–2 5–1 5–2 6–1 6–2 7–1 7–2 8–1 8–2 9–1 9–2 10–1 10–2 11–1 11–2 12–1 12–2 13–1 13–2
Sample (Patient-ROI)
g
Pearson Correlation based on Immune Cell Frequencies
Pearson R
-1
1
4-1
6-2
6-1
13-1
13-2
1-2
1-1
5-1
4-2
5-2
3-1
3-2
7-1
11-1
2-2
2-1
7-2
10-2
10-1
11-2
8-2
8-1
12-2
12-1
9-2
9-1
Sample (Patient-ROI)
h
Immune Cell Composition-based Clustering
% Variance Explained
100
75
50
25
0
1 2 3 4 5 6 7 8 9 10
### of Clusters
real data
random data
i
Chi-square Test for Cell Type Associations
adj. p value < 0.05
adj. p value ≥ 0.05
11b/c⁺ 206⁺ Mac
CD11c⁺ DC/Mac
CD14⁺ Mono
CD14⁺/16⁺ Mono
CD163⁺ Mac
CD206⁺ Mac
CD209⁺ DC
CD4 T cell
CD68⁺ Mac
CD8 T cell
γδ T cell
Giant Cell
Other Immune
Mast Cell
Neutrophil
Treg
Endothelial
Epithelial
Fibroblast
B cell
11b/c⁺ 206⁺ Mac
CD11c⁺ DC/Mac
CD14⁺ Mono
CD14⁺/16⁺ Mono
CD163⁺ Mac
CD206⁺ Mac
CD209⁺ DC
CD4 T cell
CD68⁺ Mac
CD8 T cell
γδ T cell
Giant Cell
Other Immune
Mast Cell
Neutrophil
Treg
Endothelial
Epithelial

**Extended Data Figure 2. Single cell phenotypic composition of human tuberculosis granulomas. (a)** Conceptual overview of hierarchical FlowSOM algorithm application. **(b)** Heatmap of cell lineages clustered by mean normalized protein expression of markers shown along columns. **(c)** Cell phenotype maps for all FOVs. **(d)** Total cell counts across all FOVs sorted by descending order. **(e)** Major cell lineage composition across all FOVs. Bars represent the mean and standard error. **(f)** Major cell lineage frequency of total cells broken down by FOV. **(g)** Heatmap of Pearson correlation coefficients between all FOVs based on immune cell frequencies clustered by correlation coefficient with hierarchical clustering (distance = 1 – correlation, complete linkage). **(h)** Percent variance explained per clusters based on clustering in g. Real clustering output is plotted in teal, while grey shows the results for a randomized model of immune cell frequencies. **(i)** Chi-square analysis of pairwise cell prevalence. P-values were determined with a chi-square test followed by FDR correction (FDR = 0.05).

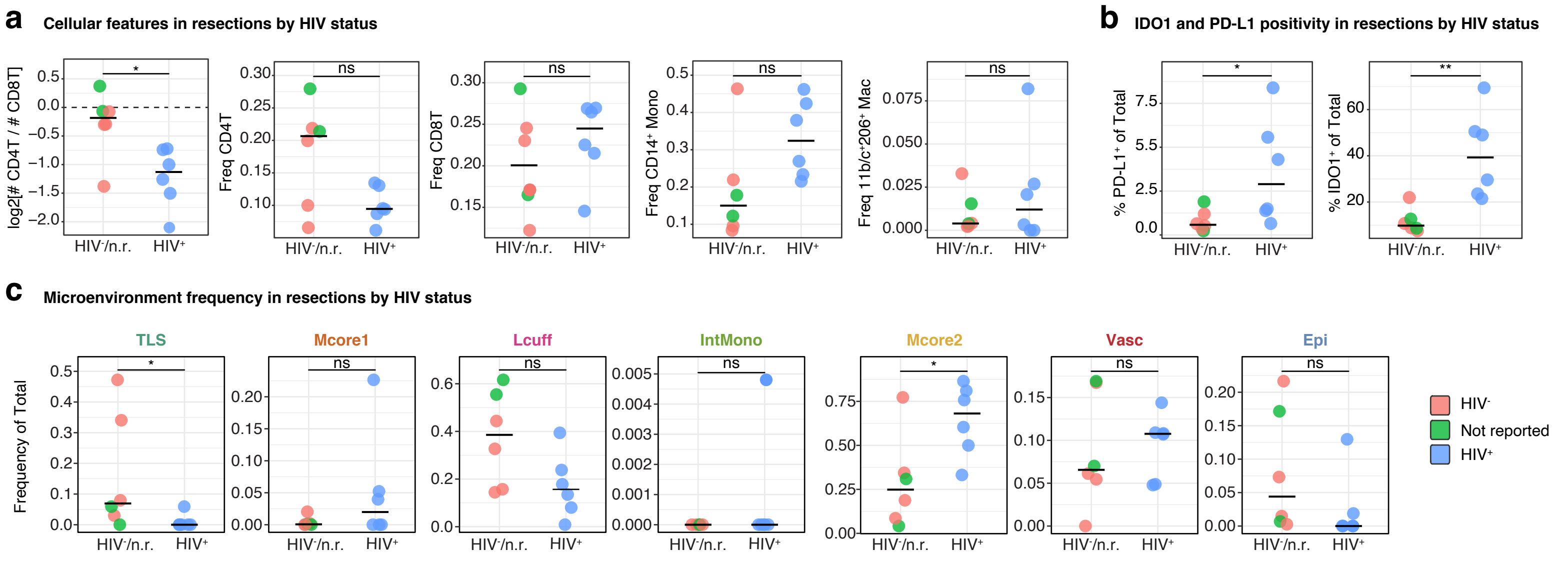
a
Cellular features in resections by HIV status
log2[# CD4T / # CD8T]
Freq CD4T
Freq CD8T
Freq CD14+ Mono
Freq 11b/c+206+ Mac
b
IDO1 and PD-L1 positivity in resections by HIV status
% PD-L1+ of Total
% IDO1+ of Total
c
Microenvironment frequency in resections by HIV status
TLS
Mcore1
Lcuff
IntMono
Mcore2
Vasc
Epi
Frequency of Total
HIV-/n.r.
HIV+
*
**
ns
HIV-
Not reported
HIV+

**Extended Data Figure 3. Analysis of cellular features of therapeutic resections with respect to HIV status. (a)** From left to right: $CD4^+$ T: $CD8^+$ T cell ratio represented as a log2 fold-change for each resection FOV, frequency of $CD4^+$ T, $CD8^+$ T cells, $CD14^+$ monocytes, and 11b/$c^+$ $206^+$ macrophages of total immune cells. Data is grouped and colored by HIV status (pink = $HIV^-$, green = not reported, blue = $HIV^+$). Line represents the median. **(b)** Frequency of $IDO1^+$ or PD-$L1^+$ cells (of total cells) across resection groups. Line represents the median. **(c)** ME frequency (of total cells) across resection groups. Line represents the median. All p-values determined with a Wilcoxon Rank Sum Test where: ns $p > 0.05$, * $p < 0.05$, ** $p < 0.01$.

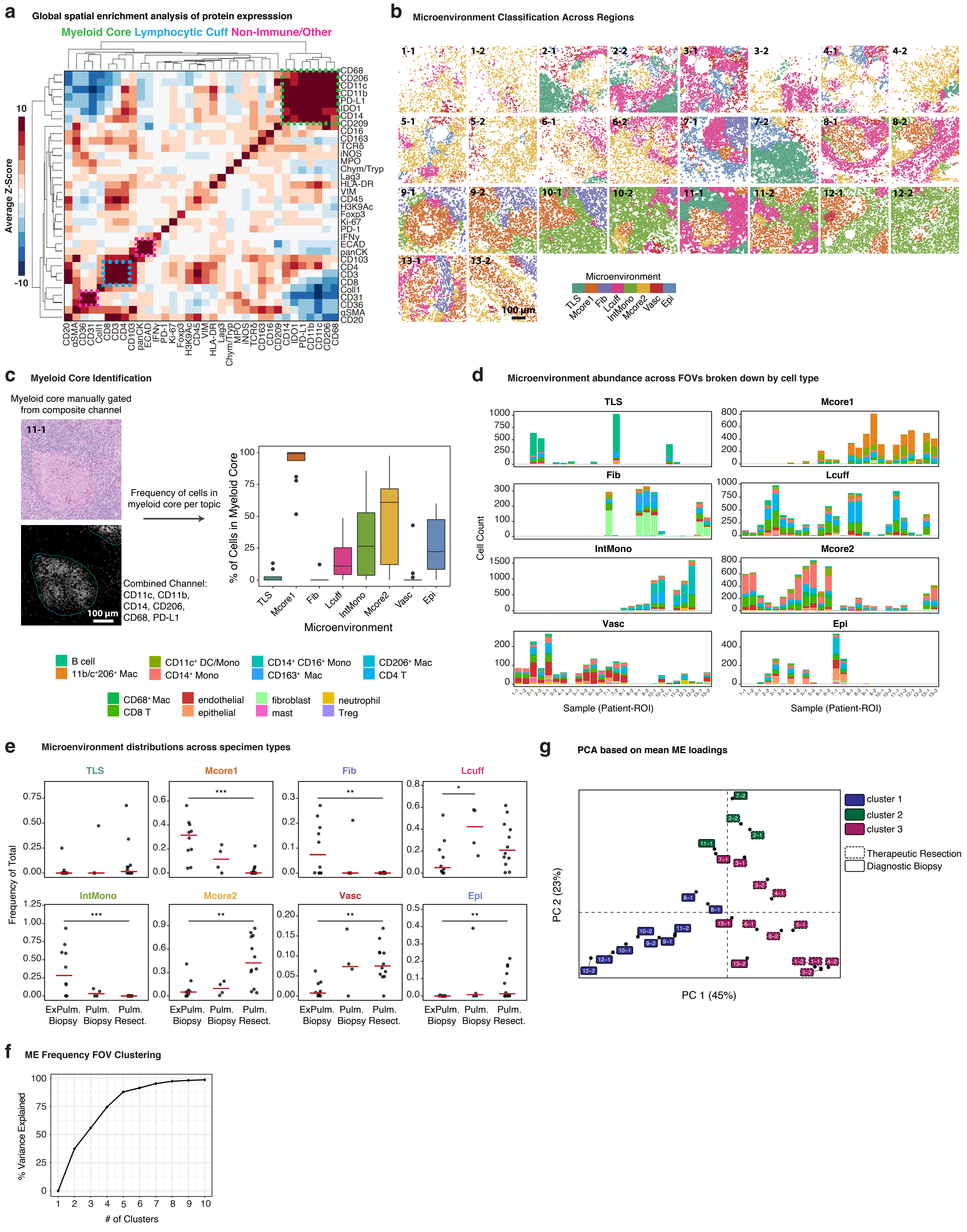

a
Global spatial enrichment analysis of protein expresssion
Myeloid Core Lymphocytic Cuff Non-Immune/Other
Average Z-Score
10
-10
b
Microenvironment Classification Across Regions
Microenvironment
TLS Mcore1 Fib Lcuff IntMono Mcore2 Vasc Epi
100 µm
c
Myeloid Core Identification
Myeloid core manually gated from composite channel
11-1
Frequency of cells in myeloid core per topic
Combined Channel: CD11c, CD11b, CD14, CD206, CD68, PD-L1
100 µm
% of Cells in Myeloid Core
Microenvironment
d
Microenvironment abundance across FOVs broken down by cell type
TLS
Mcore1
Fib
Lcuff
IntMono
Mcore2
Vasc
Epi
Cell Count
Sample (Patient-ROI)
B cell
11b/c+206+ Mac
CD11c+ DC/Mono
CD14+ Mono
CD14+ CD16+ Mono
CD163+ Mac
CD206+ Mac
CD4 T
CD68+ Mac
CD8 T
endothelial
epithelial
fibroblast
mast
neutrophil
Treg
e
Microenvironment distributions across specimen types
TLS
Mcore1
Fib
Lcuff
IntMono
Mcore2
Vasc
Epi
Frequency of Total
ExPulm. Biopsy
Pulm. Biopsy
Pulm. Resect.
g
PCA based on mean ME loadings
cluster 1
cluster 2
cluster 3
Therapeutic Resection
Diagnostic Biopsy
PC 2 (23%)
PC 1 (45%)
f
ME Frequency FOV Clustering
% Variance Explained
### of Clusters

**Extended Data Figure 4. Spatial protein enrichment and microenvironment modeling of human tuberculosis granulomas. (a)** Spatial enrichments of protein expression averaged across all TB granuloma FOVs and visualized as a heatmap hierarchically clustered (Euclidean distance, average linkage). Dashed boxes correspond to modules of protein enrichment corresponding to the myeloid core (green), lymphocytic cuff (blue), and a non-immune/other niche (pink). **(b)** Max probability maps (MaxPM) for all FOVs. **(c)** Representative hematoxylin & eosin and combined myeloid channel of a pleural TB FOV for identification of the myeloid core (left) and frequency of cells in the myeloid core across microenvironments (right). **(d)** Counts of cells broken down by phenotype across all FOVs and microenvironments. **(e)** Frequency of cells across microenvironments broken down by specimen type. **(f)** Percent variance explained per clusters based on clustering in Fig. 2f. **(g)** First two principal components of a principal component analysis (PCA) of all FOVs based on mean microenvironment probabilities. FOVs are colored by clusters from Fig. 1e and annotated by origin (therapeutic resection = dashed box, diagnostic biopsy = solid box). All boxplots represent the median and interquartile range. All p-values determined with a Wilcoxon Rank Sum Test where: ns $p > 0.05$, * $p < 0.05$, ** $p < 0.01$, *** $p < 0.001$.

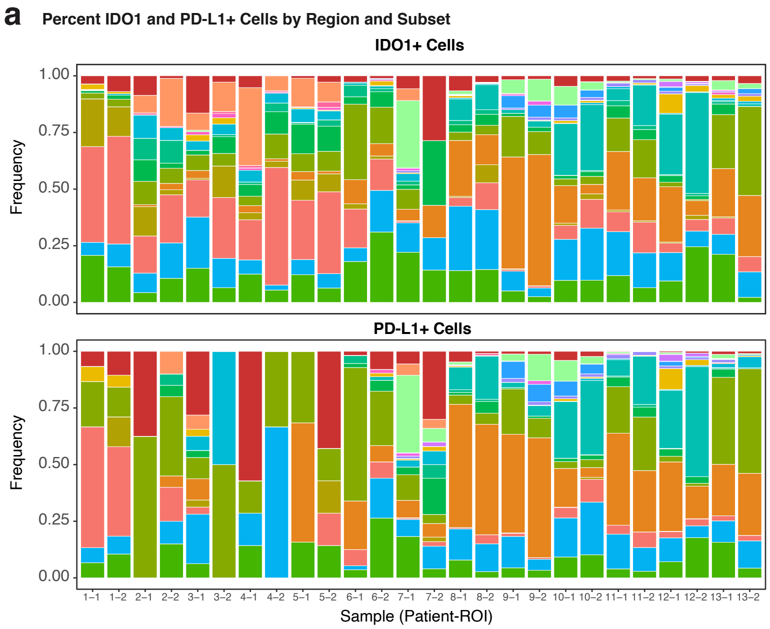


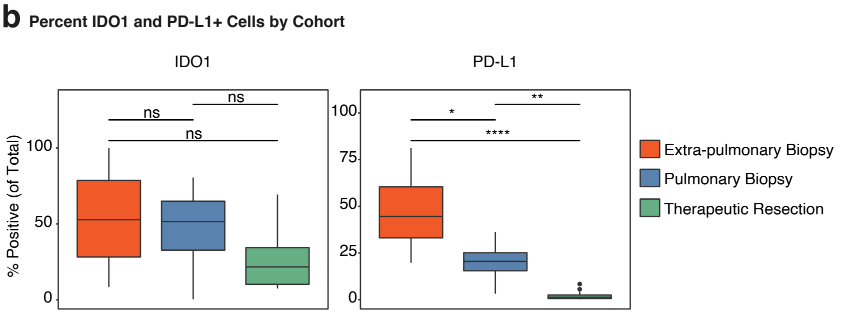


**c** Correlation between IDO1 and PD-L1 by cohort

| Cohort | Pearson R | p-value |
|---|---|---|
| Pulmonary (all) | 0.40 | p < 2.2e-6 |
| Extra-Pulmonary | 0.64 | p < 2.2e-6 |
| Resection | 0.22 | p < 2.2e-6 |
| Biopsy | 0.64 | p < 2.2e-6 |

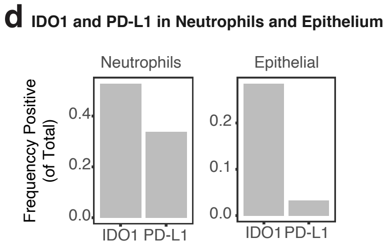


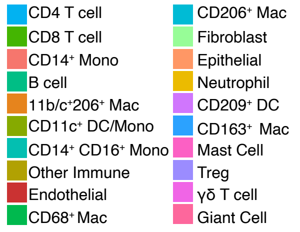


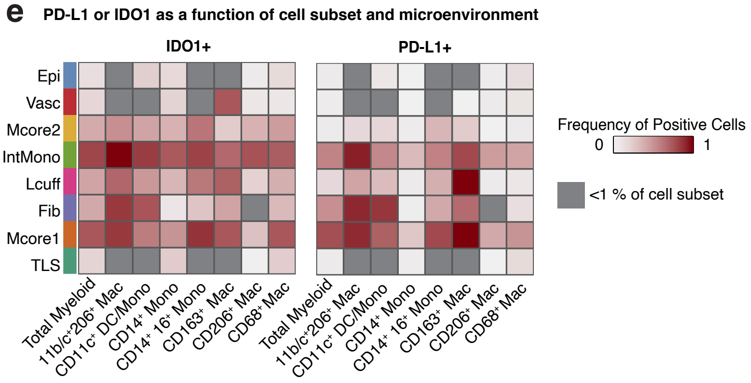


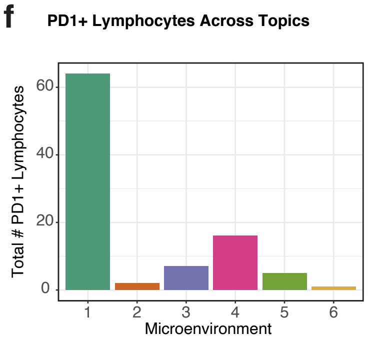

**Extended Data Figure 5. Immunoregulatory protein expression in TB granulomas. (a)** Frequency of cells positive for IDO1 (top) or PD-L1 (bottom) broken down by FOV and cell phenotype. **(b)** Frequency of $IDO1^+$ or $PD\text{-}L1^+$ cells (of total cells) across specimen type (extrapulmonary biopsy = orange, pulmonary biopsy = blue, therapeutic resection = green). **(c)** Pearson correlation coefficient and p-value determined by t-test broken down by specimen type. **(d)** Frequency of neutrophils (left) and epithelial cells (right) positive for PD-L1 or IDO1 across all FOVs. **(e)** The frequency of $IDO1^+$ and $PD\text{-}L1^+$ myeloid cells for all myeloid cells subsets across ME. Any ME with fewer than 1% of the total cell subset is shaded gray. **(f)** Frequency of $PD\text{-}1^+$ cells across all FOVs broken down by microenvironment. All boxplots represent the median and interquartile range. Unless otherwise specified all p-values were determined with a Wilcoxon Rank Sum Test where: ns $p > 0.05$, * $p < 0.05$, ** $p < 0.01$, *** $p < 0.001$.

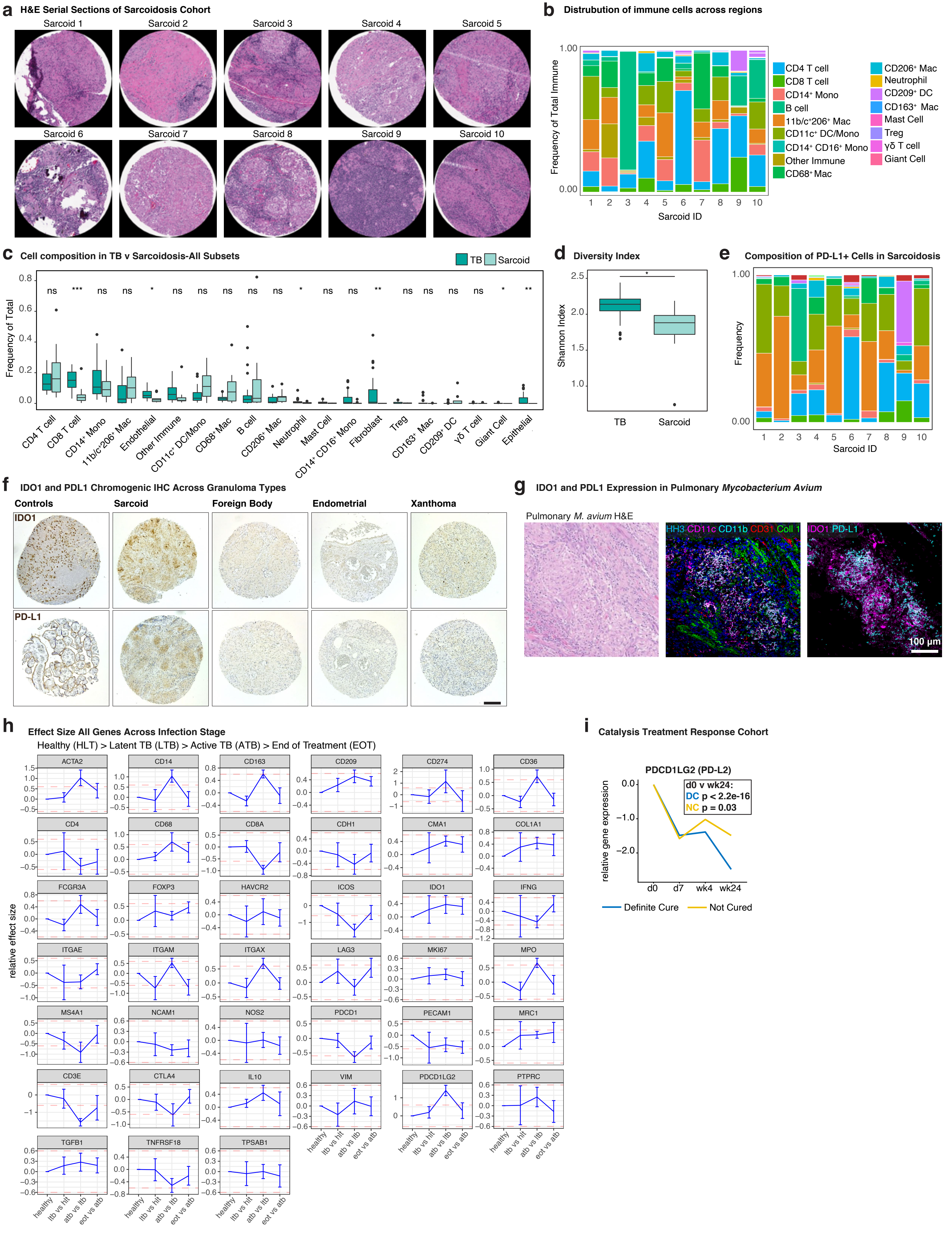
a
H&E Serial Sections of Sarcoidosis Cohort
Sarcoid 1
Sarcoid 2
Sarcoid 3
Sarcoid 4
Sarcoid 5
Sarcoid 6
Sarcoid 7
Sarcoid 8
Sarcoid 9
Sarcoid 10
b
Distrubution of immune cells across regions
Frequency of Total Immune
Sarcoid ID
CD4 T cell
CD8 T cell
CD14⁺ Mono
B cell
11b/c⁺206⁺ Mac
CD11c⁺ DC/Mono
CD14⁺ CD16⁺ Mono
Other Immune
CD68⁺ Mac
CD206⁺ Mac
Neutrophil
CD209⁺ DC
CD163⁺ Mac
Mast Cell
Treg
γδ T cell
Giant Cell
c
Cell composition in TB v Sarcoidosis-All Subsets
TB
Sarcoid
Frequency of Total
CD4 T cell
CD8 T cell
CD14⁺ Mono
11b/c⁺206⁺ Mac
Endothelial
Other Immune
CD11c⁺ DC/Mono
CD68⁺ Mac
B cell
CD206⁺ Mac
Neutrophil
Mast Cell
CD14⁺ CD16⁺ Mono
Fibroblast
Treg
CD163⁺ Mac
CD209⁺ DC
γδ T cell
Giant Cell
Epithelial
d
Diversity Index
Shannon Index
TB
Sarcoid
e
Composition of PD-L1+ Cells in Sarcoidosis
Frequency
Sarcoid ID
f
IDO1 and PDL1 Chromogenic IHC Across Granuloma Types
Controls
Sarcoid
Foreign Body
Endometrial
Xanthoma
IDO1
PD-L1
g
IDO1 and PDL1 Expression in Pulmonary Mycobacterium Avium
Pulmonary M. avium H&E
HH3 CD11c CD11b CD31 Coll 1
IDO1 PD-L1
100 µm
h
Effect Size All Genes Across Infection Stage
Healthy (HLT) > Latent TB (LTB) > Active TB (ATB) > End of Treatment (EOT)
relative effect size
ACTA2
CD14
CD163
CD209
CD274
CD36
CD4
CD68
CD8A
CDH1
CMA1
COL1A1
FCGR3A
FOXP3
HAVCR2
ICOS
IDO1
IFNG
ITGAE
ITGAM
ITGAX
LAG3
MKI67
MPO
MS4A1
NCAM1
NOS2
PDCD1
PECAM1
MRC1
CD3E
CTLA4
IL10
VIM
PDCD1LG2
PTPRC
TGFB1
TNFRSF18
TPSAB1
healthy
ltb vs hlt
atb vs ltb
eot vs atb
i
Catalysis Treatment Response Cohort
PDCD1LG2 (PD-L2)
relative gene expression
d0 v wk24:
DC p < 2.2e-16
NC p = 0.03
d0
d7
wk4
wk24
Definite Cure
Not Cured

**Extended Data Figure 6. Immunoregulatory protein expression in non-tuberculous granulomas and transcript expression in peripheral blood of TB patients. (a)** Hematoxylin & eosin stained sections of sarcoidosis granuloma FOVs. **(b)** Frequency of immune cell subsets out of total immune cells broken down by sarcoidosis FOV. **(c)** Comparison of cell type frequency (out of total cells) between tuberculosis (dark green) and sarcoidosis (light green). **(d)** Shannon diversity index in tuberculosis and sarcoidosis FOVs. **(e)** Frequency of PD-L1$^+$ cells across all sarcoidosis FOVs broken down by cell subset. **(f)** Representative immunohistochemistry images of PD-L1 or IDO1 (brown) of controls (top=spleen, bottom=placenta), a sarcoid granuloma, xanthoma granuloma, foreign body lesion, and endometrial lesion with hematoxylin nuclear counterstaining (purple). **(f)** Hematoxylin and eosin (left) and MIBI-TOF staining for major cell lineage markers (middle) or IDO1 (magenta) and PD-L1 (cyan) (right) of a representative pulmonary *Mycobacterium avium* FOV. **(h)** Gene effect sizes in in latent TB (n = 173) versus healthy controls (n = 197), latent TB (n = 372) versus active TB (n = 479), and active TB (n = 168) versus end-of-treatment (n = 160). Bars represent the mean and standard deviation. Dashed red lines represent a relative effect size of 0.6. **(i)** PD-L2 gene expression across treatment time broken down by cure status (blue = definite cure and yellow = no cure). Line represents mean expression in each time point, connected across time points. P-value determined with Student's T-test for PD-L2 expression at d0 versus wk24 in the definite cure (DC, n = 71) and not-cured (NC, n =7) groups. All boxplots represent the median and interquartile range. Unless otherwise specified all p-values were determined with a Wilcoxon Rank Sum Test where: ns $p > 0.05$, * $p < 0.05$, ** $p < 0.01$, *** $p < 0.001$.
