## Extended Data Table 1 for "Multiplexed imaging of human tuberculosis granulomas uncovers immunoregulatory features conserved across tissue and blood"

**Extended Data Table 1. Tuberculosis Granuloma Cohort Clinical Information**

**Surgical Resections**

| Patient ID | Internal ID | Soruce | Country | Specimen | Tissue | Sex | Age (yrs) | TB Type | HIV Status | Treatment | Macro/Microscopic Features |
|---|---|---|---|---|---|---|---|---|---|---|---|
| 1 | 1 | Albert Luthuli Central Hospital | South Africa | Resection | Lung | F | 35 | TB | Positive | Efavirenz/Lamivudine/Zidovudine | Right lung pneumonectomy, irregular tubercles identified, extensive granulomatous inflammation with acid-fast bacilli |
| 2 | 2 | Albert Luthuli Central Hospital | South Africa | Resection | Lung | M | 30 | TB | Negative | NA | Left lung pneumonectomy, no visible tubercles, extensive fibrosis with bronchiectasis and associated hemorrhage |
| 3 | 3 | Albert Luthuli Central Hospital | South Africa | Resection | Lung | M | 67 | MDR-TB | Not Reported | NA | Left upper lobectomy, lung contains large areas of necrotizing granulomatous inflammation, acid-fast bacilli are present |
| 4 | 4 | Albert Luthuli Central Hospital | South Africa | Resection | Lung | M | 19 | TB | Negative | NA | Left lung pneumonectomy, bronchiectatic with multiple irregularly shaped tubercles, granulomas are composed of central caseative type necrosis, paucibacillary |
| 5 | 11 | Albert Luthuli Central Hospital | South Africa | Resection | Lung | F | 37 | TB | Positive | Efavirenz/Lamivudine/Zidovudine | Right lung pneumonectomy, contains cavity with hemorrhage and necrotic material, lung parenchyma demonstrates necrotic granulomatous inflammation, paucibacillary |
| 6 | 34 | Albert Luthuli Central Hospital | South Africa | Resection | Lung | M | 24 | MDR-TB | Positive | Efavirenz/Lamivudine/Zidovudine | Left lung pneumonectomy, destroyed lung with bronchiectasis, acid-fast bacilli were identified within tubercles |

**Diagnostic Tissues**

| Patient ID | Internal ID | Soruce | Country | Specimen | Tissue | Sex | Age (yrs) | AFB | PCR |
|---|---|---|---|---|---|---|---|---|---|
| 7 | 17 | Stanford Hosptial | United States | Biopsy | Lung | F | 75 | Positive | Positive |
| 8 | 30 | Stanford Hosptial | United States | Biopsy | Lung | M | not reported | Positive | Positive |
| 9 | 18 | Stanford Hosptial | United States | Biopsy | Pleura | M | 88 | Positive | Positive |
| 10 | 20 | Stanford Hosptial | United States | Biopsy | Pleura | M | 62 | Positive | Positive |
| 11 | 31 | Stanford Hosptial | United States | Biopsy | Pleura | M | 64 | Positive | Positive |
| 12 | 29 | Stanford Hosptial | United States | Biopsy | Endometrium | F | 65 | Positive | Positive |
| 13 | 12 | Stanford Hosptial | United States | Biopsy | Lymph Node | M | 56 | Positive | Positive |
