## Extended Data Table 2 for "Multiplexed imaging of human tuberculosis granulomas uncovers immunoregulatory features conserved across tissue and blood"

**Extended Data Table 2. Multiplexed imaging antibody panel staining conditions and low-level processing parameters**

**Panel 1 (overnight stain)**

| | | | | | | | **Parameters for analysis** | | | | | |
|---|---|---|---|---|---|---|---|---|---|---|---|---|
| **Antibody target** | **Provider** | **Catalog Number** | **Lot** | **Clone** | **Mass channel** | **Titer ($\mu$g/mL)** | **Start** | **Stop** | **NoiseT 1** | **NoiseT 2** | **NoiseT 3** | **AggFilter** |
| Collagen-1 | Abcam | ab215969 | GR296572-1 | EPR7785 | 141Pr | 0.50 | 140.7 | 141.2 | 5 | 5 | 4 | 150 |
| Lag3 | LSBio | LS-C18692 | 113549 | 17B4 | 142Nd | 0.50 | 141.7 | 142.2 | 5 | 5 | 5 | 200 |
| CD4 | Abcam | ab181724 | GR32155375-1 | EPR6855 | 143Nd | 0.50 | 142.7 | 143.2 | 6 | 6 | 5 | 100 |
| CD14 | Cell Signaling | 56082BF | 2 | D7A2T | 144Nd | 0.50 | 143.7 | 144.2 | 4 | 4 | 4 | 100 |
| Foxp3 | BD Biosciences | 624084 | 8099783 | 236A/E7 | 146Nd | 1.00 | 145.7 | 146.2 | 5 | 6 | 5 | 200 |
| PD1 | Cell Signaling | 86163BF | 2 | D4W2J | 147Sm | 1.00 | 146.7 | 147.2 | 6 | 6 | 4 | 100 |
| CD31 | Abcam | ab207091 | GR241753-2 | EP3095 | 148Nd | 0.50 | 147.7 | 148.2 | 3.5 | 3.5 | 3.5 | 100 |
| PD-L1-biotin | Cell Signaling | 13684BF | 2 | E1L3N | NA | 1.00 | NA | NA | NA | NA | NA | NA |
| E-Cadherin | Abcam | ab213606 | not recorded | EP700Y | 150Nd | 0.25 | 149.7 | 150.2 | 5 | 5 | 3.5 | 180 |
| Ki67 | Cell Signaling | 9449BF | 2 | 8D5 | 151Eu | 0.25 | 150.7 | 151.2 | 4 | 4 | 4 | 100 |
| CD209/DC-SIGN | BD Biosciences | 624084 | not recorded | DCN46 | 152Sm | 0.13 | 151.7 | 152.2 | 4.5 | 4.5 | 3.5 | 150 |
| CD206 | Cell Signaling | 91992BF | 2 | E2L9N | 153Eu | 0.50 | 152.7 | 153.2 | 5 | 5 | 4.5 | 140 |
| TCRδ | Santa Cruz | sc-100289X | H2317 | H-41 | 154Sm | 1.00 | 153.7 | 154.2 | 4 | 4 | 3.5 | 200 |
| iNOS | Spring Bioscience | M4264 | 170802 | SP126 | 155Gd | 0.50 | 154.7 | 155.2 | 5 | 5 | 3 | 200 |
| CD68 | Abcam | 76437BF | 2 | D4B9C | 156Gd | 0.13 | 155.7 | 156.2 | 5 | 5 | 4.5 | 120 |
| CD36 | Abcam | 14347BF | 2 | D8L9T | 157Gd | 0.50 | 156.7 | 157.2 | 4.5 | 4.5 | 4 | 100 |
| CD8 | Cell Marque | 108M-OEM1404 | 1514101 | C8/144B | 158Gd | 0.25 | 157.7 | 158.2 | 5 | 5 | 5 | 75 |
| CD3ε | Cell Signaling | 85061BF | 4 | D7A6E | 159Tb | 0.25 | 158.7 | 159.2 | 6 | 6 | 5 | 100 |
| IDO1 | Spring Bioscience | M5604.C | 170215 | SP260 | 160Gd | 0.50 | 159.7 | 160.2 | 7 | 7 | 4.5 | 100 |
| CD11c | Abcam | ab216655 | GR3210349-1 | EP1347Y | 161Dy | 0.25 | 160.7 | 161.2 | 5 | 5 | 5 | 100 |
| CD163 | Cell Signaling | 93498BF | 2 | D5U1J | 163Dy | 2.00 | 162.7 | 163.2 | 4 | 4 | 5 | 100 |
| CD20 | Cell Marque | 120M-OEM1404 | 1429304 | L26 | 164Er | 0.50 | 163.7 | 164.2 | 4 | 4 | 5 | 100 |
| CD16 | Cell Signaling | 24326BF | 2 | D1N9L | 165Ho | 1.00 | 164.7 | 165.2 | 5 | 5 | 4 | 100 |
| IFNγ | Abcam | ab218890 | GR3191590-2 | IFNG/466 | 166Er | 1.00 | 165.7 | 166.2 | 5 | 5 | 4 | 200 |
| HLA-DR-DQ-DP | Abcam | ab7856 | GR3191247-1 | CR3/43 | 167Er | 0.25 | 166.7 | 167.2 | 5 | 5 | 4.5 | 100 |
| CD11b | Abcam | ab187537 | GR286344-1 | EP1345Y | 168Er | 0.25 | 167.7 | 168.2 | 5 | 5 | 4.5 | 100 |
| CD45 | Cell Signaling | 13917BF | 2 | D9M8I | 169Tm | 0.50 | 168.7 | 169.2 | 5 | 5 | 5 | 200 |
| H3K9Ac | Cell Signaling | 9649BF | 12 | C5B11 | 170Er | 1.00 | 169.7 | 170.2 | 3 | 3 | 3 | 200 |
| Keratin (pan) | ThermoFisher | MS-343-PABX | 343X1801A | AE1/AE3 | 171Yb | 1.00 | 170.7 | 171.2 | 4 | 4 | 4 | 150 |
| CD103 | Abcam | ab221210 | GR3175670-1 | EPR4166(2) | 172Yb | 0.50 | 171.7 | 172.2 | 5 | 5 | 4.5 | 200 |
| MPO | R&D Systems | AF3667 | YBZ0217091 | polyclonal | 174Yb | 0.75 | 173.7 | 174.2 | 3 | 3 | 3 | 180 |
| N+/K+ATPase | Abcam | ab167390 | GR3229163-1 | EP1845Y | 175Lu | 1.00 | 174.7 | 175.2 | 5 | 5 | 5 | 150 |
| HLA Class I | Abcam | ab70328 | GR307795-2 | EMR8-5 | 176Yb | 1.00 | 175.7 | 176.2 | 4 | 4 | 4 | 100 |

**Panel 2 (1h stain)**

| **Label** | **Provider** | **Catalog Number** | **Lot** | **Clone** | **Mass channel** | **Titer** | **Start** | **Stop** | **NoiseT 1** | **NoiseT 2** | **NoiseT 3** | **AggFilter** |
|---|---|---|---|---|---|---|---|---|---|---|---|---|
| HH3 | Cell Signaling | 4499BF | 7 | D1H2 | 89Y | 2.00 | 88,7 | 89.2 | 2.5 | 2.5 | 2 | 0 |
| Vimentin | Cell Signaling | 5741BF | 3 | D2H13 | 113In | 2.00 | 112.7 | 113.2 | 3 | 3 | 3 | 150 |
| SMA | Spring Bioscience | M4714.C | not recorded | SP171 | 115In | 2.00 | 114.7 | 115.2 | 2.5 | 2.5 | 2 | 200 |
| biotin | Biolegend | 409002 | B232547 | 1D4-C5 | 149Sm | 2.00 | 148.7 | 149.2 | 5 | 5 | 5 | 75 |
| Chymase | Abcam | ab233729 | GR3218665-1 | EPR13136 | 173Yb | 0.25 | 172.7 | 173.2 | 1.7 | 1.7 | 2 | 200 |
| Tryptase | Abcam | ab212156 | GR273336-1 | EPR9522 | 173Yb | 0.25 | 172.7 | 173.2 | 1.7 | 1.7 | 2 | 200 |
