## Extended Data Table 3 for "Multiplexed imaging of human tuberculosis granulomas uncovers immunoregulatory features conserved across tissue and blood"

**Extended Data Table 3. Gene Expression Cohort Description**

| Cohort | Analysis |
|---|---|
| GSE19491 | atb v hlt |
| GSE19491 | atb v ltb |
| GSE19491 | ltb v hlt |
| GSE28623 | atb v hlt |
| GSE28623 | atb v ltb |
| GSE28623 | ltb v hlt |
| GSE29536 | atb v hlt |
| GSE31348 | atb v eot |
| GSE34608 | atb v hlt |
| GSE36238 | atb v eot |
| GSE37250 | atb v ltb |
| GSE39939 | atb v ltb |
| GSE39940 | atb v ltb |
| GSE41055 | atb v hlt |
| GSE41055 | atb v ltb |
| GSE41055 | ltb v hlt |
| GSE42834 | atb v hlt |
| GSE54992 | atb v eot |
| GSE56153 | atb v eot |
| GSE62147 | atb v eot |
| GSE62525 | atb v hlt |
| GSE62525 | atb v ltb |
| GSE62525 | ltb v hlt |
| GSE73408 | atb v ltb |
| GSE74092 | atb v hlt |
| GSE74092 | atb v ltb |
| GSE74092 | ltb v hlt |
| GSE81746 | atb v hlt |
| GSE83456 | atb v hlt |
| GSE83892 | atb v hlt |
| GSE84076 | atb v eot |
| GSE101705 | atb v ltb |
| GSE107731 | atb v hlt |
| GSE119143 | atb v hlt |
| GSE40553S | atb v eot |
| GSE40553U | atb v eot |
| ACS | ACS |
| cliff | atb v eot |
| CRTC | CRTC |

| Key | |
|---|---|
| **atb** | Active TB |
| **ltb** | Latent TB |
| **eot** | End of Treatment |
| **hlt** | Healthy |
