## Extended Data Table 4 for "Multiplexed imaging of human tuberculosis granulomas uncovers immunoregulatory features conserved across tissue and blood"

**Extended Data Table 4. MetaIntegrator analysis of active TB versus healthy controls**

| gene | effectSize | effectSizeStandardError | effectSizePval | effectSizeFDR | tauSquared | numStudies | cochranesQ | heterogeneityPval | fisherStatUp | fisherPvalUp | fisherFDRUp | fisherStatDown | fisherPvalDown | fisherFDRDown |
|---|---|---|---|---|---|---|---|---|---|---|---|---|---|---|
| CD8A | -0.828553701 | 0.104622781 | 2.39E-15 | 9.30E-14 | 0.050389462 | 12 | 24.81600649 | 0.009698998 | 3.12536185 | 0.999999895 | 0.999999944 | 196.9139048 | 4.13E-29 | 4.03E-28 |
| CD3E | -1.381909016 | 0.198069911 | 3.02E-12 | 5.89E-11 | 0.280537326 | 12 | 45.14028777 | 4.58E-06 | 2.942933927 | 0.999999944 | 0.999999944 | 340.8219093 | 9.24E-58 | 3.60E-56 |
| PDCD1LG2 | 1.044332929 | 0.157800311 | 3.64E-11 | 4.73E-10 | 0.146295916 | 12 | 28.18500549 | 0.003032616 | 233.356776 | 3.20E-36 | 2.50E-35 | 2.296953309 | 0.999999996 | 1 |
| ITGAM | 1.076483238 | 0.199472155 | 6.79E-08 | 6.62E-07 | 0.279844153 | 12 | 42.9066826 | 1.13E-05 | 209.495939 | 1.50E-31 | 9.76E-31 | 1.536194882 | 1 | 1 |
| CD36 | 0.859949312 | 0.161090208 | 9.38E-08 | 7.32E-07 | 0.204122494 | 12 | 74.47020154 | 1.71E-11 | 265.6288021 | 1.29E-42 | 2.52E-41 | 10.85051706 | 0.990038697 | 1 |
| ICOS | -0.928726858 | 0.193702989 | 1.63E-06 | 1.06E-05 | 0.259041834 | 12 | 39.74938398 | 3.95E-05 | 7.288554416 | 0.999587926 | 0.999999944 | 243.1268341 | 3.79E-38 | 4.92E-37 |
| CD163 | 0.783635553 | 0.174681187 | 7.25E-06 | 4.04E-05 | 0.214685686 | 12 | 69.33669906 | 1.63E-10 | 187.8349229 | 2.32E-27 | 1.29E-26 | 3.765164186 | 0.999999265 | 1 |
| FCGR3A | 0.487532628 | 0.110433085 | 1.01E-05 | 4.59E-05 | 0.046595953 | 11 | 19.06669791 | 0.03942365 | 112.9571255 | 3.26E-14 | 1.06E-13 | 5.2625988 | 0.99990383 | 1 |
| ACTA2 | 0.920528437 | 0.209849427 | 1.15E-05 | 4.59E-05 | 0.348243226 | 12 | 50.74051454 | 4.60E-07 | 247.9199664 | 4.26E-39 | 4.16E-38 | 6.561176703 | 0.999838028 | 1 |
| CD4 | -0.692535265 | 0.158039365 | 1.18E-05 | 4.59E-05 | 0.158641546 | 12 | 31.16042266 | 0.001038737 | 8.790840639 | 0.998009236 | 0.999999944 | 139.2649438 | 3.18E-18 | 2.07E-17 |
| MPO | 0.695404051 | 0.1660425 | 2.81E-05 | 9.97E-05 | 0.168766155 | 12 | 31.08074596 | 0.001069484 | 153.4568664 | 7.53E-21 | 3.67E-20 | 7.379232178 | 0.999541125 | 1 |
| IDO1 | 0.774866873 | 0.212420832 | 0.000264504 | 0.000859639 | 0.262704777 | 10 | 31.84657789 | 0.00021166 | 138.7573995 | 8.71E-20 | 3.77E-19 | 5.783079405 | 0.999160323 | 1 |
| CD14 | 0.981011221 | 0.274968029 | 0.00036009 | 0.001080269 | 0.654378571 | 12 | 129.6267266 | 0 | 257.9593359 | 4.34E-41 | 5.65E-40 | 12.12134564 | 0.97850626 | 1 |
| TNFRSF18 | -0.500727959 | 0.162266968 | 0.002029862 | 0.005654615 | 0.172756093 | 12 | 45.93125783 | 3.33E-06 | 12.66440441 | 0.971335513 | 0.999999944 | 105.2758829 | 3.73E-12 | 1.82E-11 |
| CD274 | 1.27655224 | 0.421062805 | 0.002431499 | 0.006321897 | 2.092833485 | 13 | 262.7184236 | 0 | 460.8364537 | 4.20E-81 | 1.64E-79 | 106.9036286 | 8.89E-12 | 3.85E-11 |
| IL10 | 0.30632555 | 0.115112571 | 0.007788685 | 0.01898492 | 0.079745264 | 12 | 40.61123656 | 2.81E-05 | 90.91544911 | 1.02E-09 | 2.83E-09 | 22.40562025 | 0.555070627 | 0.746474291 |
| MS4A1 | -0.509330842 | 0.197420216 | 0.009881961 | 0.022670382 | 0.365156472 | 13 | 118.1340154 | 0 | 57.67296161 | 0.000341825 | 0.00074062 | 272.2138207 | 7.17E-43 | 1.40E-41 |
| LAG3 | -0.386130023 | 0.167998956 | 0.02153862 | 0.04666701 | 0.18223159 | 12 | 33.63106147 | 0.000415497 | 14.48675115 | 0.934882201 | 0.999999944 | 88.07223987 | 3.00E-09 | 1.06E-08 |
| PTPRC | 0.379757846 | 0.185934411 | 0.041108988 | 0.082908725 | 0.327913949 | 12 | 202.430536 | 0 | 116.7526827 | 3.66E-14 | 1.10E-13 | 58.43681219 | 0.000105817 | 0.000294775 |
| MRC1 | 0.23122355 | 0.114230994 | 0.042952125 | 0.082908725 | 0.044795317 | 9 | 16.04445161 | 0.041748274 | 52.89490043 | 2.72E-05 | 6.64E-05 | 13.89600224 | 0.735845132 | 0.956598672 |
| PDCD1 | -0.412772466 | 0.20556385 | 0.04464316 | 0.082908725 | 0.310588128 | 11 | 48.9355691 | 4.19E-07 | 16.01436202 | 0.815172355 | 0.963385511 | 104.5407348 | 1.03E-12 | 5.73E-12 |
| NCAM1 | -0.187047872 | 0.094483367 | 0.04773826 | 0.084626916 | 0.052456397 | 12 | 33.48110851 | 0.000439532 | 31.32922614 | 0.144515061 | 0.194347841 | 84.27278616 | 1.25E-08 | 4.07E-08 |
| ITGAX | 0.314735494 | 0.191326066 | 0.099965377 | 0.169506508 | 0.302992435 | 12 | 105.0064573 | 0 | 146.4917225 | 1.48E-19 | 5.78E-19 | 50.5031268 | 0.001221365 | 0.002801955 |
| CD68 | 0.361589531 | 0.227779972 | 0.11241055 | 0.182667143 | 0.495781001 | 13 | 156.7075368 | 0 | 143.9906471 | 2.62E-18 | 9.28E-18 | 38.6108758 | 0.053086371 | 0.098588974 |
| CTLA4 | -0.294867517 | 0.188358779 | 0.117475857 | 0.183262337 | 0.320445396 | 13 | 107.8652589 | 0 | 41.48339134 | 0.027728545 | 0.041592818 | 152.3602194 | 7.77E-20 | 6.06E-19 |
| CMA1 | 0.137685745 | 0.116622496 | 0.237757487 | 0.35663623 | 0.050395872 | 12 | 17.41654922 | 0.096143803 | 39.79795957 | 0.022479723 | 0.035068368 | 23.59756723 | 0.484791945 | 0.700255031 |
| TGFB1 | -0.462048879 | 0.416421165 | 0.267183903 | 0.385932304 | 1.850128268 | 12 | 391.363167 | 0 | 45.48093334 | 0.005107633 | 0.009054441 | 84.05633166 | 1.36E-08 | 4.07E-08 |
| NOS2 | 0.118174792 | 0.111329204 | 0.288467457 | 0.401793958 | 0.053310225 | 10 | 20.82502366 | 0.013450411 | 43.88373597 | 0.001559817 | 0.002896803 | 38.05223588 | 0.008726349 | 0.017911979 |
| MKI67 | 0.079805247 | 0.078660695 | 0.310320146 | 0.417327092 | 0 | 12 | 8.707830797 | 0.648840545 | 33.62306053 | 0.091594441 | 0.127577972 | 23.01013616 | 0.519199779 | 0.723171121 |
| CD209 | 0.097565064 | 0.143576559 | 0.496799965 | 0.645839954 | 0.13423952 | 12 | 31.90395772 | 0.000790266 | 57.66973162 | 0.000135214 | 0.000310196 | 33.79607962 | 0.088358992 | 0.149826116 |
| ITGAE | -0.122975443 | 0.193952668 | 0.526049021 | 0.661803607 | 0.266397944 | 12 | 43.73135699 | 8.10E-06 | 41.13889116 | 0.016081832 | 0.026132977 | 54.1447239 | 0.000407703 | 0.001055898 |
| HAVCR2 | 0.106970504 | 0.220447632 | 0.627504518 | 0.764771131 | 0.410797002 | 12 | 77.98623747 | 3.61E-12 | 89.91516762 | 1.49E-09 | 3.87E-09 | 53.94734189 | 0.000433189 | 0.001055898 |
| PECAM1 | -0.098341355 | 0.231922209 | 0.671545881 | 0.793645132 | 0.471477769 | 12 | 70.06389603 | 1.19E-10 | 38.33624397 | 0.032019106 | 0.04624982 | 94.25844327 | 2.81E-10 | 1.10E-09 |
| VIM | 0.049650097 | 0.162760816 | 0.760328493 | 0.847240404 | 0.183816756 | 12 | 53.61638631 | 1.39E-07 | 51.49080928 | 0.000911202 | 0.001776844 | 40.22567594 | 0.020223404 | 0.039435639 |
| TPSAB1 | -0.022989461 | 0.081751142 | 0.778547244 | 0.847240404 | 0.013271814 | 9 | 10.59599818 | 0.225657497 | 19.1224661 | 0.384318755 | 0.483497789 | 20.88182567 | 0.285416969 | 0.428125454 |
| COL1A1 | 0.047688602 | 0.172394404 | 0.782068066 | 0.847240404 | 0.230123208 | 12 | 70.77368141 | 8.71E-11 | 41.42530877 | 0.01495297 | 0.025355036 | 33.02098971 | 0.103631369 | 0.168400974 |
| IFNG | 0.06622974 | 0.272710923 | 0.808116224 | 0.851798182 | 0.678653888 | 12 | 172.3144383 | 0 | 52.41286824 | 0.000690976 | 0.001418319 | 48.32998209 | 0.002297006 | 0.004976847 |
| CDH1 | -0.015354139 | 0.078078344 | 0.844101116 | 0.866314303 | 0.016967086 | 12 | 15.45227358 | 0.162713459 | 23.98179729 | 0.46263863 | 0.56384083 | 35.66279813 | 0.059159563 | 0.10487377 |
| FOXP3 | -0.001586775 | 0.080386328 | 0.984251289 | 0.984251289 | 0 | 12 | 10.84236152 | 0.456562797 | 26.42461843 | 0.331977444 | 0.431570677 | 27.9828023 | 0.260765979 | 0.406794927 |
